# Unsupervised Clustering Enables Cohort-Specific Variant Recalibration in Crops

**DOI:** 10.64898/2026.10.07.757277

**Authors:** Thomas Bergmann, Tessa R. MacNish, Jacqueline Batley, David Edwards

**Affiliations:** Centre for Applied Bioinformatics, University of Western Australia, 6009 Perth, Australia; School of Biological Sciences, University of Western Australia, 6009 Perth, Australia

**Keywords:** variant discovery, variant recalibration, crop genomics, SNPs, INDELs, GATK, Gaussian mixture model, unsupervised learning, cohort-specific, Nextflow

## Abstract

Trait discovery in crops require reliable marker datasets, yet variant discovery remains constrained by the lack of standardised, high-confidence reference resources. Publicly available plant genomic resources often rely on outdated reference genomes, heterogenous pipelines, and generic variant hard-filtering thresholds derived from human studies. Consequently, applying the Genome Analysis Toolkit (GATK) Best Practices framework, particularly machine learning-based variant quality score recalibration (VQSR), remains challenging in crops lacking suitable training resources.

Here, we present an unsupervised approach for variant quality assessment and generating cohort-specific training resources, enabling integration of VQSR into established GATK workflows without predefined high-confidence training variant sets. Using *Brassica napus* as a model, we use the Illumina 60k *Brassica* SNP array to establish a reference-specific truth set for base quality score recalibration (BQSR) and independent benchmarking. Gaussian mixture model (GMM)-based clustering of multidimensional variant annotation profiles identifies high-, intermediate-, and low-quality variant classes, which provide flexible training resources without requiring labelled training data.

We further validate the approach in diploid *Brassica oleracea* and hexaploid wheat (*Triticum aestivum*), representing contrasting population characteristics and genome architectures. Across all three species, GMM-based clustering consistently identifies informative quality classes and enables cohort-specific recalibration of both SNPs and insertion-deletions (INDELs). Comparison with conventional hard filtering reveals discordant pass/fail classifications for up to 46.4% of SNPs and 37.8% of INDELs across evaluated cohorts, demonstrating that the approaches capture distinct aspects of variant quality. Our results establish unsupervised variant classification as transferable strategy for generating cohort-specific VQSR resources, extending established GATK workflows to crops lacking comprehensive genomic resources.

## Introduction

Genome-wide marker datasets are now widely used as the primary input for trait discovery and genomic selection in plant breeding. Genomic prediction models use genome-wide variation to estimate breeding values and accelerate genetic gain (Alemu et al. 2024; Meuwissen et al. 2001). Because predictive performance depends strongly on the quality of the underlying genotype data (Alemu et al. 2024), generating high-confidence marker datasets is essential to minimise technical artefacts and systematic errors that can propagate into downstream analyses and ultimately influence breeding decisions.

GATK (DePristo et al. 2011; McKenna et al. 2010) has become a widely established framework for genetic variation discovery, particularly in human genomics (The 1000 Genomes Project Consortium et al. 2015; Karczewski et al. 2020; Chen et al. 2024). Well-documented protocols (Van der Auwera and O’Connor 2020; Van Der Auwera et al. 2013) and comprehensive databases containing billions of curated genetic variants from large-scale human cohorts, including gnomAD (Karczewski et al. 2020) or dbSNP (Phan et al. 2025), support reliable and harmonised variant discovery.

Although GATK has been widely applied for variant discovery in plant genomics (Wu et al. 2019a; Hu et al. 2022; Wang et al. 2018; Zhou et al. 2024), often outperforming other variant calling tools (Wu et al. 2019b; Schilbert et al. 2020; Yao et al. 2020), the implementation of its Best Practices workflows remains challenging in many crop species. In particular, VQSR, which uses machine-learning models to assess variants based on multiple annotation metrics, relies on suitable high-confidence training and truth resources that are often unavailable for crops (Van der Auwera and O’Connor 2020; Van Der Auwera et al. 2013). Consequently, variant discovery in plant genomics often relies on generic hard-filtering approaches that apply fixed thresholds to individual annotations. Though practical, these approaches do not model relationships among multiple quality metrics and may not adequately capture cohort-specific variation in variant quality.

Reference variant sites serve two distinct purposes in the GATK Best Practices workflow: BQSR during data pre-processing and VQSR during variant filtering (Van Der Auwera et al. 2013). VQSR uses reference high-confidence variants to train a GMM that learns variant context annotation profiles associated with high- and low-quality variants in the cohort. The resulting variant quality score (VQSLOD) is then used to apply sensitivity-based filtering thresholds across the call set. This approach, commonly referred to as soft filtering, enables a flexible filtering according to the desired balance between variant sensitivity and precision.

When suitable reference variant resources are unavailable, variant filtering commonly relies on hard filtering. This approach retains or removes variants according to predefined thresholds for individual annotation metrics. Although simple to implement, hard filtering is less powerful than machine learning-based approaches, requires profound analytical understanding of the variant annotation distributions, and may discard true variants that fail a single filtering criterion (Van der Auwera and O’Connor 2020). Consequently, it is not considered part of the GATK Best Practices framework (Van der Auwera and O’Connor 2020; Van Der Auwera et al. 2013). Due to the lack of reference variant resources, hard filtering remains widely used in crop genomics. However, the filtering thresholds recommended by GATK, which were developed and evaluated using human genomic datasets (Van der Auwera and O’Connor 2020), are frequently adopted in plant studies without systematic assessment of their suitability for different crop species, genome architectures, and population characteristics (Lu et al. 2019; Yan et al. 2021; Cheng et al. 2024).

Applying GATK Best Practices to crop genomics is further complicated by the availability of multiple reference genomes within species, extensive genetic diversity, and complex genome architectures. Although numerous genomic databases have been developed for crop species (Thudi et al. 2021), their suitability as high-confidence reference resources for GATK workflows is often limited. For example, several databases for *Brassica* provide extensive collections of genomic variation (Yan et al. 2021, 2020; Yang et al. 2022, 2023; Scheben et al. 2019), but many rely on outdated reference genomes (Yan et al. 2021, 2020; Yang et al. 2023), lack accessible variant call files (Scheben et al. 2019), or were generated using variant-calling and filtering strategies that differ substantially from GATK Best Practices (Yan et al. 2021, 2020; Yang et al. 2022). As a result, these databases may not provide suitable reference variant sets for BQSR or the training and truth resources required for VQSR, limiting the implementation of these recalibration steps in crop-genomics workflows.

Here, we present an unsupervised clustering approach for variant quality assessment and the construction of cohort-specific training resources for VQSR, enabling its integration into established GATK workflows for crop species lacking comprehensive reference variant resources. Using tetraploid *Brassica napus* as a model system, we utilise the Illumina 60k *Brassica* SNP array (Clarke et al. 2016) to establish a reference-specific truth set for BQSR and independently evaluate an unsupervised machine-learning approach for constructing VQSR resources. We demonstrate that GMM-based clustering of multidimensional variant annotation profiles identifies high-, intermediate-, and low-quality variant classes, which can be used to generate cohort-specific VQSR training resources without requiring predefined high-confidence training variants.

We further evaluate the framework across populations of diploid *Brassica oleracea* and hexaploid wheat (*Triticum aestivum*), demonstrating that cohort-specific GMM-derived resources support VQSR of both SNPs and INDELs across species, population sizes, and levels of genetic diversity. Although SNP array data are used for independent benchmarking in *Brassica* and wheat, the approach does not depend on array-derived truth sets or predefined high-confidence training variants. It can therefore be applied to a broad range of crop species, including those with established SNP array resources, such as rice (Zhang et al. 2023), as well as species for which such resources are unavailable.

## Results

### Overview of the unsupervised variant discovery workflow

Our variant discovery framework is implemented as a scalable Nextflow pipeline, complemented by R scripts for statistical analysis and visual exploration of variant annotation profiles, cluster assignments, and model performance. The workflow follows the GATK Best Practices (DePristo et al. 2011; Van Der Auwera et al. 2013), with two key modifications. First, where available, variant sites derived from SNP arrays are used as known-sites resources for BQSR (Figure 1A) and to define a reference-specific truth set for independent evaluation of variant quality profiles. When array-derived resources are unavailable, BQSR is initially omitted, and the workflow can be repeated using high-confidence variants identified in the initial VQSR round as known sites for BQSR, following the iterative strategy described in the GATK Best Practices (Van Der Auwera et al. 2013). In addition to correcting systematic sequencing-related biases in base quality estimates, BQSR can help mitigate differences in quality-score calibration between sequencing datasets. This is particularly relevant for data sets in which base quality scores have been binned differently as pointed out in this study.

**Figure 1.**
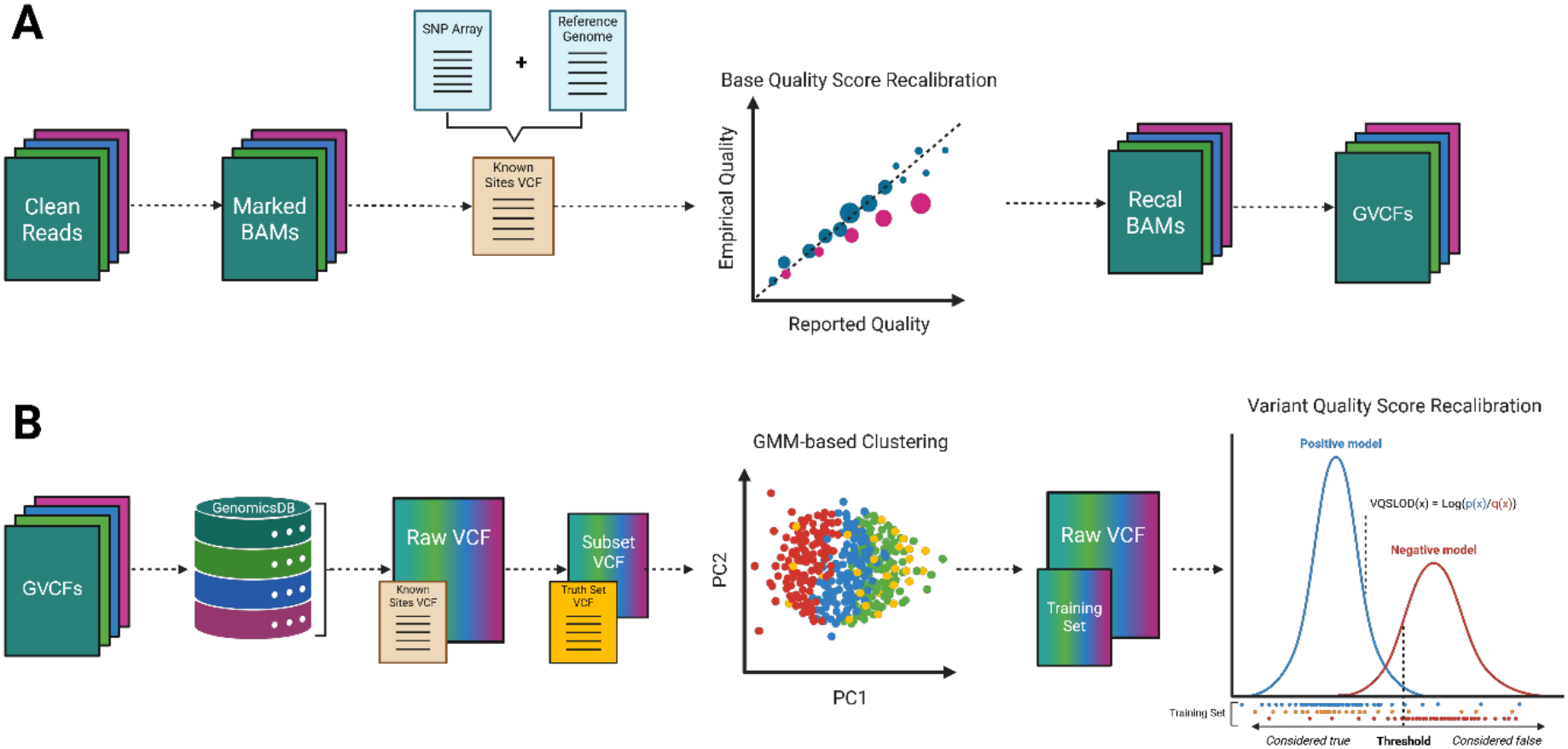
Overview of the workflow for cohort-specific variant recalibration in crop genomes. **A** Following read alignment and duplicate marking, reference-specific known variant sites derived from SNP arrays, such as the Illumina 60k *Brassica* SNP array, are used for base quality score recalibration (BQSR). For species lacking SNP array resources, BQSR can initially be omitted and subsequently performed using high-confidence variants identified in an initial round of variant discovery. Recalibrated BAM files are then processed using GATK Haplotype Caller in reference-confidence (ERC GVCF) mode. **B** Joint genotyping of GVCFs produces a cohort-level VCF. Representative subsets of SNPs and INDELs are independently clustered using Gaussian mixture models (GMMs) based on their multidimensional variant annotation profiles, assigning variants to high-and intermediate-confidence classes. The two classes are used to construct cohort-specific training resources for variant quality score recalibration (VQSR), which is subsequently applied to the full variant call set. Where available, SNP array-derived truth sets are used independently to evaluate clustering performance. Illustrations are adapted from Van der Auwera and O’Connor (2020) and created with Biorender.

Second, we introduce a GMM-based clustering step between joint genotyping and VQSR, in which representative subsets are randomly selected from the cohort-wide call set and SNPs and INDELs are clustered independently to generate cohort-specific training resources (Figure 1B). This clustering step is based on a systematic evaluation of multidimensional variant annotation profiles and identifies high-, intermediate-, and low-quality classes without requiring predefined high-confidence training sets. Where available, annotation distributions from SNP array-derived truth sets provide an independent reference for characterizing population-specific profiles associated with high-confidence variants and evaluating the quality classes identified by GMM clustering. The high- and intermediate-quality clusters are subsequently used as training resources for VQSR, allowing the resulting recalibration model to be applied to the full cohort-wide call set. By considering relationships among multiple annotation metrics rather than relying on individual, predefined filtering thresholds, this approach provides a flexible strategy for variant quality assessment and filtering that can be tailored to the characteristics and requirements of individual cohorts.

### SNP array-derived variant sites enable base quality score recalibration in *B. napus*

By mapping and filtering the Illumina 60k *Brassica* SNP array against the *B. napus* reference genome ‘Darmor-bzh’ (Rousseau-Gueutin et al. 2020), we identified 36,628 high-confidence known variant sites distributed across the genome, with an average inter-SNP distance of 23.4 kb (Supplementary Data 1, Table S1). Incorporating these known sites into BQSR substantially reduces systematic sequencing biases across all evaluated covariates, including reported base quality score, dinucleotide context, and machine cycle. For example, in sample CRR379121, the root mean square error (RMSE) decreases from 10.5 to 1.65 for base quality score bias, from 7.77 to 0.43 for dinucleotide context, and from 8.46 to 0.48 for machine cycle effects (Figure 2). Comparable reductions are observed in additional randomly selected samples from the *B. napus* mini-core collection (Supplementary Data 1, Figures S1-S3).

**Figure 2.**
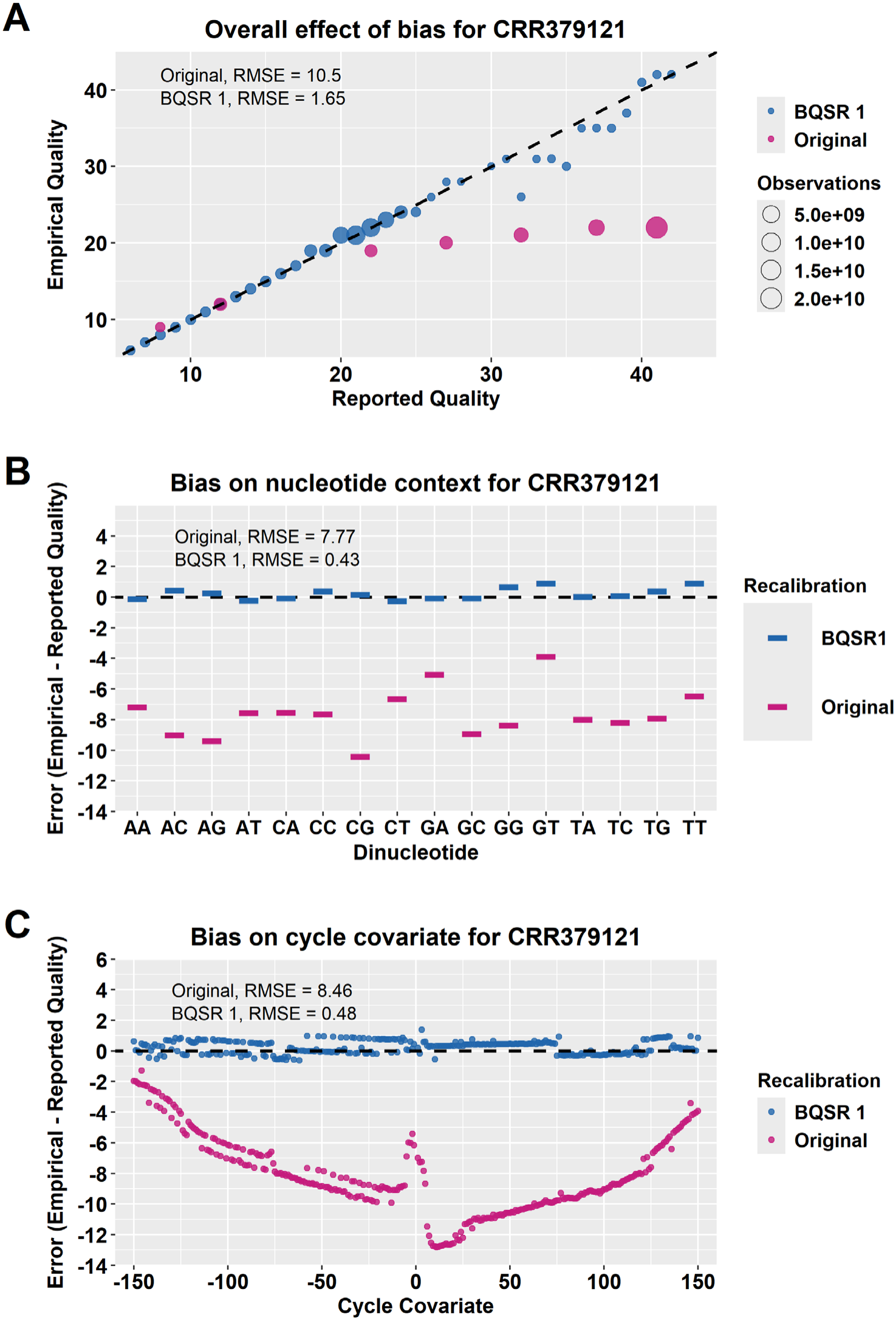
Base quality score recalibration (BQSR) with the 36k known sites in the Darmor-bzh reference genome (v. 10) for **A** base quality score bias, **B** dinucleotide context bias, and **C** machine cycle covariate bias in sample CRR379121 from the *Brassica napus* mini-core collection. Pink = uncalibrated data; blue = recalibrated data. The raw uncalibrated data contains base quality scores in seven bins: Q8, Q12, Q22, Q27, Q32, Q37, and Q41.

### Constructing a high-confidence variant set using SNP array-derived sites

Following variant calling, SNP array-derived known variant sites are matched against the unfiltered cohort VCF to establish a cohort-specific truth set. This identifies 34,784 high-confidence sites in the *B. napus* mini-core collection, hereafter referred to as the 34k truth set. We use this set as an independent benchmark for evaluating the unsupervised clustering approach. The 34k truth set exhibits a transition-to-transversion (Ti/Tv) ratio of 2.3, compared with 1.4 in the unfiltered SNP call set across the entire cohort (Supplementary Data 1, Table S2). To construct a representative dataset for subsequent clustering analyses, we then randomly select one million SNPs from the cohort VCF and characterize their genomic distribution across the Darmor-bzh reference genome. The selected subset shows broad genome-wide coverage, with an average inter-SNP distance of 846 bp (Supplementary Data 1, Table S3).

### SNP array-derived variants are characterized by high-quality annotation profiles

Comparing variant annotation profiles between the 34k truth set and the one-million-SNP subset from the *B. napus* mini-core collection reveals a clear enrichment of high-quality variants in the SNP array-derived truth set. The QualByDepth (QD) distribution of the one-million-SNP subset shows a distinct peak of low-quality variants around QD ≈ 2, alongside two peaks at higher values around QD ≈ 25 and QD ≈ 33 (Supplementary Data 1, Figure S4A). In contrast, the 34k truth set lacks the low-quality QD peak and is enriched with higher QD values, exhibiting a major peak around QD ≈ 25 followed by a plateau around QD ≈ 33. Similar enrichment for high-quality annotation values is observed across the other evaluated metrics, including Fisher Strand (FS), Strand Odds Ratio (SOR), Mapping Quality (MQ), MQRankSum (MQRS), and ReadPosRankSum (RPRS), further supporting the high-confidence nature of the SNP array-derived truth set (Supplementary Data 1, Figure S4B-F). Consistent with these annotation profiles, approximately 99% of truth-set variants pass conventional hard filtering thresholds, compared with 64% of variants in the one-million-SNP subset. The hard-filtering pass rate in the subset is comparable to that of the complete cohort call set, supporting its representativeness for subsequent analyses.

### Annotation space in the *B. napus* cohort is primarily structured by strand bias

To characterize the multidimensional annotation space used for clustering, we perform principal component analysis (PCA) on SNP annotation profiles from the *B. napus* cohort. Strand bias metrics (FS and SOR) contribute most strongly to the first principal component (PC1), which explains approximately 37% of the total variance. PCs 2-4 each explain 15-18% of the variance, whereas subsequent components account for less than 10% each (Supplementary Data 1, Figure S5-6). The dominant contribution of strand bias persists when either FS or SOR is removed from the analysis. Although FS and SOR are moderately correlated (*R^2^* = 0.53, *p < 2.2 x 10^-16^*), they quantify strand bias using different statistical approaches, indicating that their contribution to the annotation structure is not simply attributable to redundancy between the two metrics. Together, these results reveal structured variation across multiple annotation dimensions, providing a basis for unsupervised GMM clustering to identify groups of variants with distinct quality profiles.

### Unsupervised GMM clustering identifies high-quality variant classes

We evaluate GMMs containing up to ten Gaussian components and select the model that provides the best balance between model complexity and goodness of fit based on the Bayesian Information Criterion (BIC). The model selection identifies a three-component GMM as the best-fitting solution (Figure 3A), using only multidimensional variant annotation profiles and without requiring prior variant labels or external truth sets. The selected model differentiates SNPs into high-, intermediate-, and low-quality annotation profiles (Table 1). Independent evaluation against the 34k truth shows that the high-quality cluster closely matches the characteristic annotation profile of truth-set variants, including high QD and MQ values, low strand-bias metrics, and balanced mapping- and read-position metrics. Of the truth-set variants represented in the clustering subset, 82% (1,151) are assigned to the high-quality cluster, while 14% (191) and 4% (54) are assigned to the intermediate- and low-quality clusters, respectively. Hence, the unsupervised GMM identifies a high-quality annotation class that resembles the annotation profiles of the truth-set variants without requiring labelled training data. Projection of the variants into PCA space shows separation among the three clusters, with strand-bias metrics contributing strongly to the variation associated with cluster structure (Supplementary Data 1, Figure S7).

**Figure 3.**
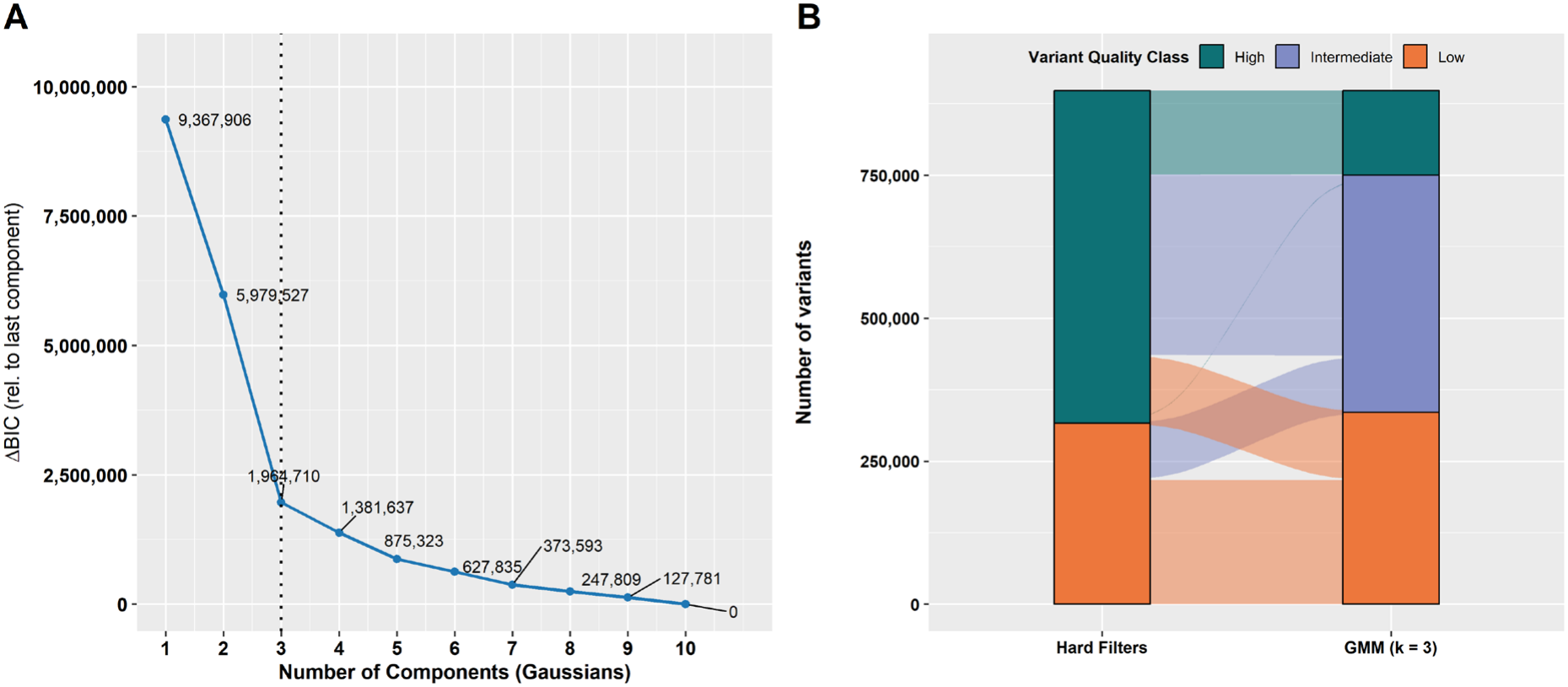
Model selection and variant quality class stratification using Gaussian mixture model (GMM)-based clustering. **A** Change in Bayesian Information Criterion (ΔBIC) relative to the largest model (*k* = 10) across increasing numbers of Gaussian components (*k*). Lower values indicate better model support. The dashed vertical line indicates best-model-fit. **B** Alluvial plot showing the flow of variant quality classification between hard filtering and GMM-based clustering (*k* = 3). Each stratum represents the classification at a given stage, and its vertical width corresponds to the number of variants. Alluvia are coloured according to group assignments from the GMM with *k* = 3.

**Table 1.** Summary statistics for SNP quality classes based on GMM clustering in the *Brassica napus* mini-core collection.

| **SNP Set** | **Variant Class** | **#SNPs** | **#True SNPs\*** | ***Med(x)*** |  |  |  |  |  |
|---|---|---|---|---|---|---|---|---|---|
|  |  |  |  | **QD** | **FS** | **SOR** | **MQ** | **MQRS** | **RPRS** |
| Clustering Subset\*\*<br>(*k* = *3*) | High | 147,814 | 1,151 | 22.6 | 0.9 | 0.7 | 60 | 0 | 0.1 |
|  | Intermediate | 414,560 | 191 | 13.5 | 1.9 | 0.8 | 54 | -1.9 | 0.15 |
|  | Low | 335,650 | 54 | 5.6 | 26.4 | 3.1 | 53 | -2.9 | 0.05 |
| 34k Truth Set | Pass | 34,441 | 34,441 | 23.4 | 0.6 | 0.7 | 60 | 0 | 0.1 |
|  | Fail | 343 | 343 | 19.2 | 48.8 | 3.5 | 56 | -1.9 | 0.12 |

*k* number of clusters*; QD* QualByDepth*; FS* FisherStrand*; SOR* StrandOddsRatio*; MQ* MappingQuality*; MQRS* MappingQualityRankSum*; RPRS* ReadPosRankSum
\*Variants overlapping with the 34k truth set
\*\*Clustering Subset with SNPs containing only complete annotation profiles

Comparing with conventional hard filtering, GMM-based clustering largely preserves the high-and low-quality variant classes while additionally identifying an intermediate-quality class comprising variants with multidimensional annotation profiles that cannot be classified using fixed, one-dimensional thresholds (Figure 3B).

### Constructing cohort-specific truth and training resources for VQSR

The three-component GMM provides a framework for constructing cohort-specific resources for VQSR. We use the high-quality cluster to define the truth resource, comprising variants with the highest-confidence annotation profiles, and the intermediate-quality cluster to construct the training resource for model recalibration. The posterior probabilities provide an additional measure of confidence in cluster assignment. More than 90% of variants are assigned to their respective cluster with posterior probabilities of ≥ 90%. Furthermore, the proportion of variants violating conventional hard-filtering thresholds increases progressively from the high- to the intermediate- and low-quality clusters, providing independent support for the quality gradient identified by GMM clustering (Supplementary Data 1, Table S4).

We notice that variants with lower posterior probabilities (25-50%) are enriched for annotation outliers across all three clusters. These variants frequently receive similar posterior probabilities across the clusters (e.g., approximately 33% for each cluster) and are assigned to the largest GMM cluster by default, which can correspond to any variant quality class. We therefore recommend a minimum posterior probability threshold of ≥50% to retain variants with more confident cluster assignments. For the *B. napus* cohort, we apply more stringent thresholds of ≥80% for the truth resource and ≥70% for the training resource, yielding 147,182 and 403,849 variants, respectively. Adjusting these thresholds allow the generation of cohort-specific VQSR resources tailored to project-specific requirements.

### Cohort-specific variant quality recalibration and tranche filtering in the *B. napus* cohort

Training the VQSR model with the high- and intermediate-quality SNP clusters enables recalibration of variant quality scores across the full call set and subsequent filtering based on truth-sensitivity tranches (Table 2). At a truth-sensitivity threshold of 99.50% -corresponding to a VQSLOD cutoff that retains 99.50% of the variants in the GMM-derived truth resource – 13,331,967 novel SNPs are retained. Diagnostic plots of the recalibration model show that VQSR captures distinct annotation profiles associated with high- and low-quality SNPs, with strand-bias metrics emerging as major features distinguishing retained from filtered variants.

**Table 2.**
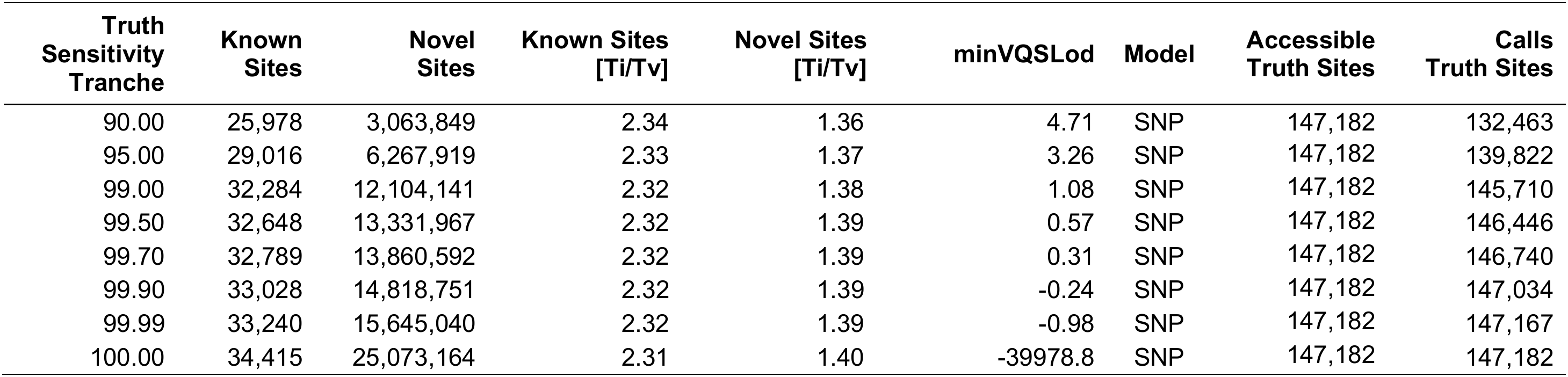
Summary statistics for cohort-specific SNP quality score recalibration across different truth sensitivity tranches in the *Brassica napus* mini-core collection.

To further evaluate the annotation profiles of recalibrated variants, we compare the distributions of retained and filtered SNPs in a second one-million-variant subset sampled from the recalibrated VCF. At the 99.50% truth-sensitivity tranche, retained variants consistently exhibit annotation profiles characteristic of high-confidence SNPs, whereas filtered variants are enriched for annotation values associated with lower-quality calls across all six evaluated metrics (Figure 4). The strongest separation is observed for the strand-bias metrics FS and SOR, consistent with their dominant contribution to the annotation structure identified by PCA.

**Figure 4.**
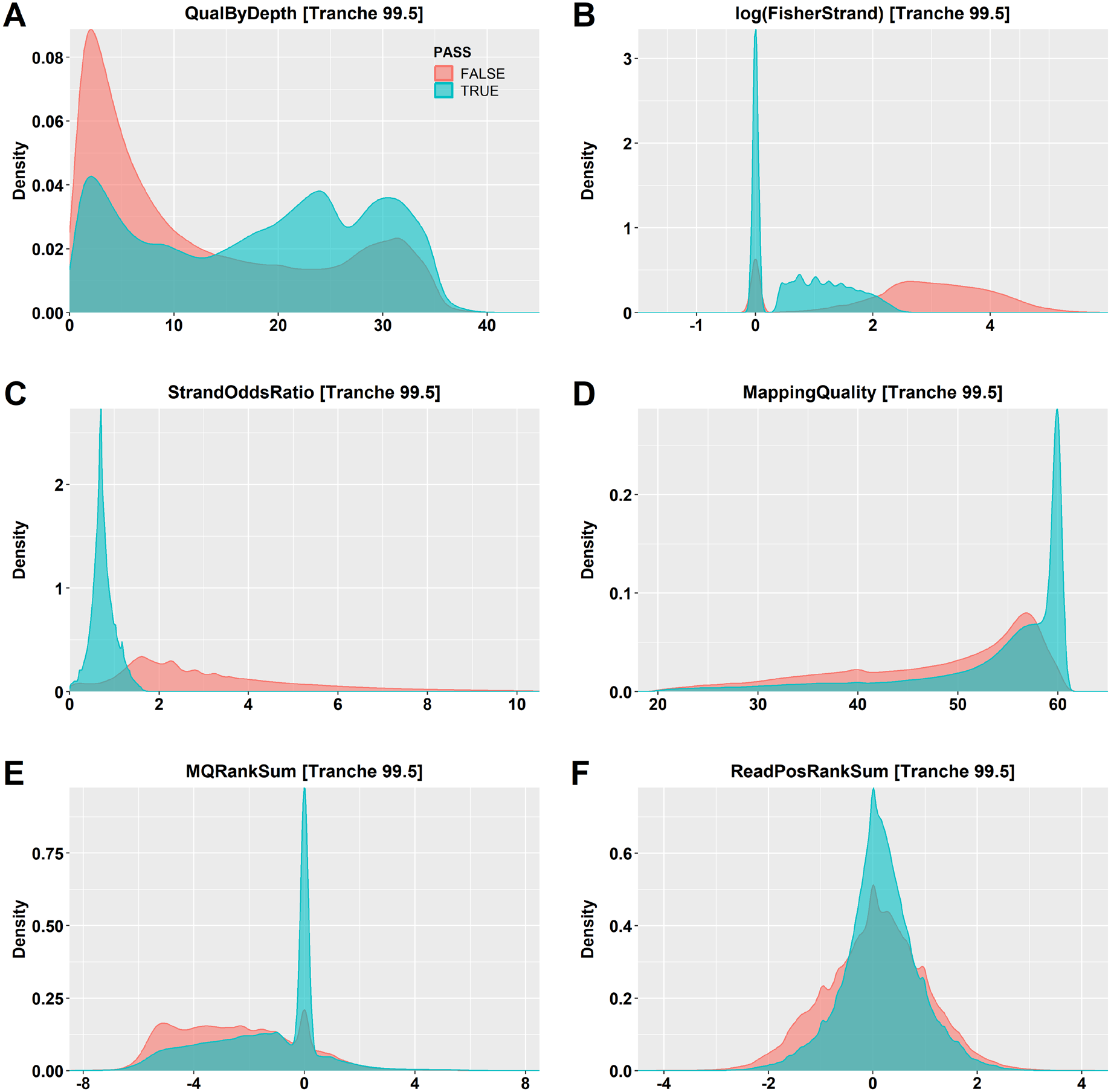
Distribution of **A** QualByDepth, **B** log-transformed Fisher Strand, **C** Strand Odds Ratio, **D** Mapping Quality, **E** MQRankSum, and **F** ReadPosRankSum values for retained (blue) and filtered (red) SNPs following variant quality score recalibration (VQSR) and filtering at the 99.5% tranche in a one-million-variant subset from the *Brassica napus* mini-core collection.

Applying the same framework to INDELs identifies a four-component GMM as the best-fitting model, distinguishing very-high, high-, intermediate-, and low-quality annotation profiles (Supplementary Data 1, Table S5). Using the very-high- and the high-quality cluster to construct the truth resource and the intermediate-quality cluster as the training resource for VQSR yields 3,098,067 recalibrated INDELs at a truth-sensitivity threshold of 99.50% (Supplementary Data 1, Table S6). As observed for SNPs, retained INDELs are enriched for high-quality annotation profiles (Supplementary Data 1, Figure S8), demonstrating applicability of the unsupervised recalibration framework applies to both major classes of small variants. Comparison of VQSR tranche filtering with conventional hard filtering reveals discordant pass/fail classifications for 28.0% of SNPs and 37.8% of INDELs in the evaluated subsets.

### SNP array resources enable BQSR across crop datasets

We next evaluate the applicability of SNP array-derived resources across crop species and populations differing in ploidy, genetic diversity, and population size. We apply the workflow to a small wild *B. oleracea* collection (n = 7, approximately 112x coverage) and a small *T. aestivum* population (n = 6, approximately 12x coverage), using genome-wide distributed high-confidence SNP sites from the Illumina 60k *Brassica* SNP array (Supplementary Data 2, Table S1) and the Wheat 660k SNP array (Supplementary Data 3, Table S1), respectively. Incorporating these sites into BQSR reduces systematic sequencing biases in base quality estimates and helps mitigate differences in quality-score calibration between the sequencing datasets (Supplementary Data 2, Figures S1-S3; Supplementary Data 3, Figures S1-S3). In both populations, the SNP array-derived sites also enable the construction of cohort-specific truth sets enriched for high-quality annotation profiles (Supplementary Data 2, Figure S4; Supplementary Data 3, Figure S4).

### Variant annotation profiles differ across crop populations

Multidimensional analysis of variant annotation profiles reveals that strand-bias metrics are major contributors to annotation variation across all evaluated datasets, although their relative contributions differ between populations. In the diploid *B. oleracea* population, both FS and SOR contribute strongly to PC1, which explains approximately 34% of the total variance (Supplementary Data 2, Figures S5-S6), consistent with the pattern observed in *B. napus*. However, the correlation between FS and SOR is weaker in *B. oleracea* (*R^2^* = 0.29, *p < 2.2 x 10^-16^*) than in *B. napus*. In the wheat dataset, FS is the dominant contributor to annotation variation, whereas SOR contributes less strongly to the principal components (Supplementary Data 3, Figures S5-S6). Consistent with this difference, FS and SOR show only a weak correlation in wheat (*R^2^* = 0.03, *p < 2.2 x 10^-16^*), indicating that the two metrics capture largely distinct patterns of variation in this population.

### Unsupervised clustering captures population-specific variant quality profiles

A three-component GMM is selected as the best-fitting model in both the *B. oleracea* and *T. aestivum* populations, distinguishing SNPs with high-, intermediate-, and low-quality annotation profiles while capturing population-specific differences in variant quality (Supplementary Data 2, Figure S7; Supplementary Data 3, Figure S7). In *B. oleracea*, annotation profiles follow a pattern similar to that observed in *B. napus*, with quality gradually decreasing across the annotation metrics from the high- to the low-quality cluster (Supplementary Data 2, Table S4). Both strand-bias metrics are positively associated with the posterior probability of assignment to the low-quality cluster (FS: *R^2^* = 0.31, *p < 2.2 x 10^-16^*; SOR: *R*^2^ = 0.51, *p < 2.2 x 10^-16^*).

A different pattern is observed in the wheat population. Here, the high- and intermediate-quality clusters differ primarily in MQ (median values of 60 and 40, respectively), whereas the low-quality cluster is characterized by lower QD (approximately 7.4) and higher strand-bias metrics (FS approximately 4.8; SOR approximately 2.2) (Supplementary Data 3, Table S4). Correlation analysis shows that higher MQ values are associated with a greater posterior probability of assignment to the high-quality cluster (*R*^2^ = 0.25, *p < 2.2 x 10^-16^*), whereas higher FS values are associated with a greater posterior probability of assignment to the low-quality cluster (*R*^2^ = 0.50, *p < 2.2 x 10^-16^*). Approximately 72% of variants in the wheat dataset have missing MQRS and RPRS values. We therefore excluded these metrics from clustering in this population.

In both populations, the majority of truth-set variants are assigned to the high-quality cluster, accounting for approximately 80% in *B. oleracea* and 57% in *T. aestivum*. Projection of the variants into PCA space further illustrates the separation of the three quality classes and highlights the substantial contribution of strand-bias metrics to cluster structure in both datasets (Supplementary Data 2, Figure S8; Supplementary Data 3, Figure S8).

### Cohort-specific clustering enables variant recalibration across crop populations

In both populations, the high-quality SNP cluster is used to define the truth resource, while the intermediate-quality cluster provides the training resource for VQSR, enabling cohort-specific variant recalibration using truth-sensitivity tranches (Supplementary Data 2, Table S5; Supplementary Data 3, Table S5). Diagnostic plots of the recalibrated models show that VQSR captures the annotation characteristics represented by the cohort-specific training and truth resources and uses these patterns to distinguish variants across the full call sets. Consistent with the clustering results, retained variants are enriched for annotation profiles associated with high-quality SNPs in both populations (Supplementary Data 2, Figure S9; Supplementary Data 3, Figure S9). Comparison of VQSR tranche filtering with conventional hard filtering across both populations reveals discordant pass/fail classifications for up to 46.4% of SNPs and 30% of INDELs in the evaluated subsets.

## Discussion

### SNP array-derived variant resources improve sequence data pre-processing

SNP array-based genotyping platforms provide cost-effective, highly reproducible genotype calls and are available for many major crop species (Rasheed et al. 2017). Their widespread use in genetic mapping, trait discovery, and breeding programs makes them an established source of high-confidence polymorphic sites in crop genomics (Sun et al. 2017; Li et al. 2014; Zhang et al. 2023; Sun et al. 2020). Incorporating array-derived variant sites into BQSR therefore provides a practical means of reducing systematic biases in base-quality estimates before variant calling, using resources that are already available for many crop species. Although sequencing technologies continue to improve, systematic errors persist (Ma et al. 2019), highlighting the relevance of BQSR in sequencing-based variant discovery.

SNP arrays typically represent only a subset of the genetic variation within a population. Nevertheless, the availability of these known polymorphic sites provides a useful basis for BQSR. In our datasets, incorporating array-derived sites reduces systematic biases in base-quality estimates, resulting in more consistent recalibration across samples. This is particularly relevant for population-scale studies in which base quality scores are differently binned between batches, as observed in the wheat cohort (Supplementary Data 3, Figure S1).

### SNP array resources are useful but not required for variant quality recalibration

An additional advantage of SNP array-derived resources is their flexibility. As demonstrated here using the Illumina *Brassica* 60k SNP array and previously for the Wheat 660k SNP array (Sun et al. 2025), array markers can be remapped to different reference genomes. Importantly, although SNP array-derived variant sites provide a useful resource for BQSR and independent evaluation, they are not required for the unsupervised GMM-based framework presented here. When suitable known-sites resources are unavailable, BQSR can be omitted during an initial round of variant discovery and performed subsequently using high-confidence sites identified from the resulting call set, following an iterative approach described in GATK Best Practices (Van Der Auwera et al. 2013).

### SNP arrays enable reference-specific construction of independent truth sets

Beyond their role in BQSR, SNP array-derived variant sites provide valuable reference-specific resources for independently evaluating variant annotation profiles. By matching array markers to the cohort-wide variant call set, high-confidence sites can be identified and their annotation profiles compared with those of randomly sampled cohort-wide variants. In our study, this comparison shows that the high-quality GMM cluster captures the characteristic annotation profiles of independently identified high-confidence variants. Importantly, the array-derived truth set is used exclusively for independent evaluation and is not required for GMM model selection or construction of the cohort-specific VQSR resources. Consequently, the framework remains applicable to variant classes for which genotyping resources are unavailable, such as INDELs, and to crop species lacking established SNP array resources or other predefined high-confidence variant sets.

### Ascertainment and mapping bias in SNP array-derived variant resources

However, SNP array-derived sets of known variant sites are subject to ascertainment bias. They are based on pre-selected, predominantly biallelic SNPs easily detectable by genotyping assays. Consequently, they underrepresent rare and population-specific variants (Rasheed et al. 2017). This bias is further intensified by our extraction strategy for the *Brassica* cohorts, in which only unambiguously aligned, full-length probe sequences are retained. Probes with multiple genomic alignments, which are particularly common in *Brassica* genomes owing to their history of whole genome triplication (Lysak et al. 2005; Chalhoub et al. 2014), are excluded to minimize uncertainty in marker position. Although this restriction improves the reliability of the extracted sites, it further limits their genomic representation and may introduce mapping-related bias into the resulting truth sets. More permissive extraction strategies could increase marker representation but would require additional validation to ensure their reliability.

In the hexaploid wheat population, MQ is an important metric distinguishing high- and intermediate-quality annotation profiles among truth-set variants. This may partly reflect the challenges of assigning sequence reads to unique genomic locations in a highly repetitive and homoeologous genome. In this study, we use only the Type I and Type II SNPs from the 660k array, which are defined by their unique best BLAST alignments to the ‘AK58’ reference genome (Sun et al. 2025). However, the presence of additional lower-scoring alignments may indicate additional sequence similarity at other genomic locations, potentially contributing to ambiguity in read mapping.

### Ti/Tv ratios are population- and species-specific

Differences between SNP array-derived known sites and novel variants are also reflected in their Ti/Tv ratios. In the *Brassica* populations, array-derived sites show Ti/Tv ratios of approximately 2.3, compared with 1.4 in the *B. napus* genome-wide SNP set and 1.52 in *B. oleracea*. Similar ratios (1.32 – 1.39) have been reported in previous *B. napus* studies (Huang et al. 2013; Bus et al. 2012; Dalton-Morgan et al. 2014), as well as in other crops, including maize (1.53) (Ching et al. 2002). Higher ratios have been reported for intergenic marker sets in *B. rapa* (approximately 1.6) (Park et al. 2010) and barley (approximately 2.39) (Rostoks et al. 2005). In contrast, the wheat cohort exhibits higher Ti/Tv ratios for both array-derived sites (approximately 2.45) and novel SNPs (approximately 1.95), highlighting species-specific differences in transition-transversion composition.

The response of Ti/Tv ratios to variant filtering also differs between populations. While the ratios of array-derived known sites remain relatively stable across truth-sensitivity thresholds, novel SNPs show population-specific changes. The Ti/Tv ratio remains largely unchanged in *B. napus*, increases with less stringent filtering in *B. oleracea*, and decreases in wheat. These contrasting patterns indicate that changes in filtering stringency can affect the Ti/Tv composition of retained variants differently across populations.

Although a Ti/Tv ratio of approximately 3.0 in exonic regions is often considered indicative of high-quality SNP call sets in human genomics (Wang et al. 2015), Ti/Tv bias varies across species (Keller et al. 2007) and is influenced by genomic sequence context (Morton 1995; Morton et al. 1997). Our results demonstrate that these ratios vary between crop species and populations and respond differently to variant filtering. Consequently, Ti/Tv thresholds derived from human genomic studies should not be transferred directly to genome-wide variant discovery in crops. Instead, they should be evaluated empirically within the target species and population.

### Filtering parameters must be adapted to crop genomes

Similar to Ti/Tv ratios, variant quality metrics should be evaluated in the context of the species, genome architecture, and population structure rather than adopted directly from human genomic studies. Commonly used hard-filtering thresholds derived from human sequencing data may not adequately account for the characteristics of complex crop genomes, such as hexaploidy wheat. In our wheat population, a substantial number of variants, including independently validated truth-set variants, exhibit MQ values around 40 while retaining annotation profiles characteristic of high-quality variants across other metrics. Applying the commonly used GATK hard-filtering threshold of MQ < 40 therefore excludes many variants that are supported by the multidimensional annotation framework presented here. This observation illustrates how reliance on individual annotation thresholds can lead to the removal of variants whose combined annotation profiles are otherwise consistent with high confidence. More broadly, variant quality annotations reflect both technical and genome-specific properties, highlighting the need to evaluate filtering criteria within the context of the target species and population. The framework presented here provides a means of assessing these properties jointly, supporting more context-aware variant quality assessment in crop genomics.

### Unsupervised GMM-based clustering enables cohort-specific VQSR

Our framework uses GMM to identify distinct variant annotation profiles and construct cohort-specific resources for VQSR. This approach takes advantage of the probabilistic assignment of GMMs, allowing variants to be assigned to clusters according to their posterior probabilities rather than through strictly discrete classifications. Compared with commonly used clustering approaches such as K-Means (Raykov et al. 2016), GMMs offer greater flexibility in modelling heterogenous, multidimensional annotation profiles. In particular, GMMs allow clusters to differ in variance and covariance structure, while the number of components can be evaluated based on model fit rather than fixed a priori (Maugis et al. 2009). Whereas K-Means assigns variants to discrete clusters assuming spherical clusters with equal variance, while also being sensitive to outliers and differences in cluster density (Raykov et al. 2016). The variation in annotation profiles observed across our datasets, including the presence of intermediate-quality variants and overlapping annotation distributions, makes the flexibility of GMMs particularly useful for distinguishing cohort-specific variant classes.

### Alternative modelling approaches may extend variant quality recalibration

Bayesian mixture models represent a potential extension of the GMM approach presented here. Such approaches could provide additional flexibility in modelling variant annotation profiles; however, whether these improve the sensitivity or robustness of unsupervised variant quality recalibration remains to be evaluated and is beyond the scope of this study. Deep learning-based approaches have emerged as alternatives to conventional variant-calling and quality assessment methods (Junjun et al. 2024; Olson et al. 2022). Tools such as DeepVariant (Poplin et al. 2018) and GATK’s CNNScoreVariants (Friedman et al. 2020) use neural networks trained on human sequencing datasets to evaluate variant quality. Although they can be applied to crop species, their performance requires careful evaluation, particularly where suitable high-confidence reference variant sets are unavailable. Even though species-specific models can be trained (Friedman et al. 2020; Yun et al. 2019), their accuracy depends strongly on the quality, size, and representativeness of the training data (Junjun et al. 2024; Yun et al. 2019).

Our results further demonstrate that variant annotation profiles differ between species and populations. In the *Brassica* populations, strand-bias metrics are major contributors to variant classification, whereas MQ is an important discriminator in wheat. Rank-sum metrics (MQRS and RPRS) are less informative in our datasets and are particularly limited in the wheat population, where approximately 72% of variants lack values for these annotations at a sequencing depth of approximately 12x. These differences highlight the potential limitations of applying variant quality models across datasets with contrasting annotation distributions, sequencing depths, and genome architectures. Consequently, models developed or trained using human genomic data should be evaluated in the target crop species and population before being adopted for variant quality assessment.

## Methods

This section describes the datasets analysed and the workflow for cohort-specific variant recalibration in crop genomes. The workflow was applied to the datasets described below, with detailed cohort reports provided in the supplementary data. The variant calling pipeline was implemented in Nextflow (Di Tommaso et al. 2017) and incorporates GATK4 version 4.6.2.0, SAMtools (Li et al. 2009) version 1.23.1, BWA (Li and Durbin 2009) version 0.7.19, and BCFtools (Danecek et al. 2021) version 1.23.1. The unsupervised clustering was conducted in R (R Core Team 2024).

### Dataset of a mini-core collection of *Brassica napus*

Raw genome sequencing data from a mini-core collection of a world-wide collection of *B. napus* accessions was obtained from the Genome Sequence Archive (GSA) database in BIG Data Centre (http://bigd.big.ac.cn/gsa) under BioProject accession number PRJCA008062 (Hu et al. 2022). The dataset comprises 135 *B. napus* accessions including 72 semi-winter, 26 spring-, and 37 winter-types. The average sequencing depth was 45x across all samples. Quality control of the raw sequencing data was performed via fastp (Chen et al. 2018) with the following parameters: *--correction --detect_adapter_for_pe*. The pseudochromosomes of the *B. napus* cv. *Darmor-bzh* (Rousseau-Gueutin et al. 2020) reference genome (v. 10) were used for read mapping and variant calling. The reference genome was split into equally sized intervals with *gatk SplitIntervals* and the following options: *--scatter 100 --subdivision-mode INTERNAL_SUBDIVISION*.

### Dataset of a wild collection of *Brassica oleracea*

Raw genome sequencing data of seven accessions of wild *B. oleracea* were obtained from the NCBI repository under BioProject accession number PRJNA947374 (Ji et al. 2024). The dataset included the following wild *Brassica oleracea* species: *B. macrocarpa*, *B. cretica*, *B. insularis*, *B. villosa*, *B. incana*, *B. oleracea* var. *oleracea*, and *B. oleracea* ‘W03’. The average sequencing depth was 112x across all seven accessions. Quality control of raw sequencing data was performed via fastp as described before. The pseudochromosomes of *B. oleracea* ‘W03’ reference genome (Ji et al. 2023) were used for read mapping and variant calling. The reference genome was prepared as described before.

### Dataset of a mini collection of *Triticum aestivum*

Raw genome sequencing data of three modern cultivars and three landraces of the A.E. Watkins landrace collection were obtained from the GSA database in BIG Data Centre under BioProject accession number PRJCA019636 (Cheng et al. 2024). The average sequencing depth was 12x across all samples. Quality control of raw sequencing data was performed via fastp as described before. The pseudochromosomes of the elite winter wheat cultivar ‘Aikang 58’ (AK58) reference genome (Jia et al. 2023) were used for read mapping and variant calling. The reference genome was prepared as described before with the modification that it was split into 200 equally sized intervals.

### Generation of reference-specific known sites from SNP arrays

Probe sequences of the Illumina 60k SNP *Brassica* SNP array were aligned against both *Brassica* reference genomes and high-confidence known SNP sites were extracted with a single in-house Bash script (see Supplementary Data 1 for details). Refined SNP positions of the Wheat 660k SNP array (Sun et al. 2025) were used to generate a reference-specific known-site resource in the ‘AK58’ genome. To reduce the amount of unambiguously mapped probe sequences, only Type I and Type II SNPs, corresponding to markers with a unique best BLAST alignment to the ‘AK58’ reference, were included and converted to a VCF file.

### Alignment and data pre-processing

Whole-genome sequencing reads were aligned to the reference genomes using *bwa mem* with default parameters. Resulting BAM files were sorted and duplicate reads were marked using *gatk MarkDuplicates*. BQSR was performed using *gatk BaseRecalibrator* and *gatk ApplyBQSR*, using known variant sites derived from the SNP arrays. The recalibration data was inspected using *gatk AnalyzeCovariates -csv*, and recalibration plots were generated using custom R scripts.

### Variant calling and joint genotyping

Variant calling was performed per sample and genome interval using *gatk HaplotypeCaller* in *ERC GVCF* mode. Per-sample interval GVCFs were consolidated into cohort-specific interval databases using *gatk GenomicsDBImport*. Joint interval genotyping was carried out with *gatk GenotypeGVCFs*, producing raw cohort-level interval VCF files. The interval VCFs were concatenated into one cohort VCF using *bcftools concat*. Summary statistics about the raw VCFs were generated with *gatk VariantEval*.

### Construction of cohort-specific training datasets

Unfiltered cohort VCFs were converted to variant annotation tables using *gatk VariantsToTable* or *bcftools query*. For the *Brassica* populations, one-million variants were randomly selected from the annotation tables and the known sites derived from the 60k *Brassica* SNP array were used to generate cohort-specific truth sets. The positions of the one-million variant subsets were plotted and checked to ensure their uniform distribution across the *Brassica* genomes. To account for the large genome size of wheat, the AK58 genome was divided into 5 kb intervals, and a single variant site was randomly sampled from each interval. The resulting dataset comprised a representative genome-wide subset of approximately three million variants distributed across the entire wheat genome. The clustering subsets and truth sets were imported into R (R Core Team 2024). Variants in the SNP clustering subsets matching the truth sets were labelled as true variants. Data manipulation was performed with the dplyr package (Wickham et al. 2023).

SNPs and INDELs were analyzed independently, and separate clustering models were fitted for each variant class. For SNPs, the following variant annotations were included in the analysis: QualByDepth (QD), FisherStrand (FS), StrandOddsRatio (SOR), MappingQuality (MQ), MappingQualityRankSumTest (MQRS), and ReadPosRanKSumTest (RPRS). For INDELs, clustering was performed without array-derived truth sets. The annotations QD, FS, SOR, MQRS, and RPRS were included in the analysis.

### Variant context annotation analysis

Within each cohort, variant context annotation distributions from the clustering subsets and, for SNPs, the array-derived truth sets were visualized to assess their characteristics and relationships. Pairwise Pearson correlation coefficients between annotation metrics were calculated using the *cor()* function in base R. Variant annotations were Z-score normalized using the *scale()* function. Variants containing missing annotations values were excluded from PCA and clustering analysis.

PCA was performed independently for SNPs and INDELs on approximately 100,000 randomly selected variants from each clustering subset using the *prcomp()* function (c*enter = FALSE, scale = FALSE*). The factoextra package (Kassambara and Mundt 2020) was used to generate scree plots via *fviz_eig*() and loading plots via *fviz_pca_var*(), enabling assessment of variance explained and annotation contributions to principal components. The resulting PCA spaces were retained as a reference framework for downstream visualization and comparison of clustering structures. Heatmaps were generated using the ComplexHeatmap package (Zuguang Gu 2017), and plots were created with ggplot2 (Wickham 2016).

### Variant quality stratification via GMM clustering

GMM-based clustering was used to identify groups of variants with similar annotation profiles. Z-score normalized variant context annotations were used to fit a GMM using the *GMM()* function from the ClusterR package (Mouselimis 2025; Sanderson and Curtin 2016). Model fitting was performed using the following parameters: *gaussian_comps = 10, dist_mode = “maha_dist”, seed_mode = “random_subset”, km_iter = 10, em_iter = 100*. The optimal balance between model complexity and goodness of fit was evaluated using differences in the Bayesian Information Criterion (ΔBIC), calculated for each model relative to the model containing 10 Gaussian components. The optimal number of components (*k*) was defined as the point where substantial reductions in ΔBIC were followed by only marginal improvements with additional components.

Clustering was performed independently for SNPs and INDELs within each cohort. The resulting clusters represented distinct variant annotation profiles that were subsequently evaluated by comparing cluster-specific annotation distributions, enrichment of true variants (where available), and enrichment of hard-filter PASS and FAIL variants.

### Evaluation of cluster annotation profiles

To evaluate clustering performance, summary statistics of variant annotations were compared across clustering approaches, and distributions were visualized using boxplots generated with the ggplot2 package (Wickham 2016). Alluvial plots (Brunson and Read 2023; Brunson 2020) were used to visualize the flow of variants and to identify core variant clusters. Cluster structure and separability were further evaluated by projecting variants onto the previously defined PCA space and visualizing them using PCA biplots generated with the factoextra package via the *fviz_pca_biplot*() function. Variants were colored according to their GMM-derived cluster assignments, enabling visual assessment of cluster separation and overlap within the annotation space.

### Cohort-specific variant quality score recalibration

The clustering procedure was designed to identify well-defined variant sets representing high-and intermediate-confidence variant calls for downstream recalibration. Variants assigned to the high-confidence cluster were used as truth resource (*known=false, training=true, truth=true, prior=15.0*). Variants assigned to the intermediate-confidence cluster were used as training resource (*known=false, training=true, truth=false, prior=12.0*). Where available, SNP array-derived variants were additionally incorporated as known-site resources (*known=true, training=false, truth=false, prior=2.0*). These resources were supplied to *gatk VariantRecalibrator* to construct independent cohort-specific recalibration models for SNPs and INDELs. Variant filtering was performed using *gatk ApplyVQSR* based on selected tranche sensitivity thresholds. The resulting VQSR models were assessed using customized versions of the diagnostic plots generated by *VariantRecalibrator*. To evaluate the recalibrated variant call sets for a given tranche sensitivity threshold, random subsets of SNPs and INDELs were sampled from the recalibrated VCFs and analyzed in R as described above.

## Data access

The data sets used in this study are publicly available. The scripts, as well as resources for the SNP array data, are available at GitHub (https://github.com/tbergmann92/Advanced-Crop-Variant-Discovery).

## Acknowledgements

This work was supported by resources provided by the Pawsey Supercomputing Centre with funding from the Australian Government and the Government of Western Australia. We acknowledge funding from the Australian Research Council (Projects FL230100030, DP200100762, DP240100051, DP210100296, IC230100016 and LP230100351) and from the Grains Research and Development Corporation (UWA2307-001RTX, UWA2307-002RTX and UWA2604-002BGX).

## Author contributions

Conceptualization was by T.B. Methodology was by T.B. Investigation was by T.B. and T.R.M. Formel analysis was by T.B. and T.R.M. Validation was by T.B. Visualization was by T.B. Writing of the original draft was by T.B. Reviewing and editing were by T.B., T.R.M., J.B., and D.E. Supervision was by J.B. and D.E. Funding acquisition was by T.B., J.B. and D.E.

