## Supplementary Data 1 for "Unsupervised Clustering Enables Cohort-Specific Variant Recalibration in Crops"

### Unsupervised Clustering Enables Cohort-Specific Variant Recalibration in Crops

Thomas Bergmann[1,2], Tessa R. MacNish[1,2], Jacqueline Batley[2], and David Edwards[1]

[1]Centre for Applied Bioinformatics, University of Western Australia, 6009 Perth, Australia

[2]School of Biological Sciences, University of Western Australia, 6009 Perth, Australia

#### Variant Calling and Filtering in a *Brassica napus* Mini-Core Collection

<u>Scripts and resources:</u>

https://github.com/tbergmann92/ Advanced-Crop-Variant-Discovery

#### Preparation of SNP array-derived known variant sites in *Brassica napus*

1. Input data

- Brassica_60k_50bp_ProbeSeqs.fasta (provided in *Resources*)
- Brassica_60k_REF_ALT_Alleles.csv (provided in *Resources*)
- DarmorV10_Chromosomes_Only.fa (not provided)

2. Preparation of reference genome

The reference genome is indexed using:

- *bwa index*
- *samtools faidx*
- *gatk CreateSequenceDictionary*

3. Generation of SNP-array derived known sites (integrated pipeline)

All downstream processing, including sequence alignment, alignment filtering, and SNP extraction, is performed using a single in-house Bash script:

- 4_Prepare_60k_SNP_Array_DarmorV10.sh

Within this script:

- probe sequences are aligned to the reference genome using *bwa aln* (parameters: *-n 0.01 -o 0 -l 32 -k 2*)
- alignments are converted to BAM format using *bwa samse*
- alignments are sorted using *samtools sort*
- high-confidence mappings are retained by filtering:
    - primary alignments only (*-F 0x904*)
    - uniquely mapped probes (*X0:i:1*)
    - exclusion of alternative alignments (*XA:Z:* absent)
    - full-length alignments (*CIGAR = 50M*)
- SNP positions are extracted directly from filtered alignments
- probe identifiers are mapped to reference and alternate alleles using the provided annotation table (Brassica_60k_REF_ALT_Alleles.csv)
- a sorted VCF file of high-confidence SNP array-derived variants is generated

The final output consists of:

- a sorted VCF file of high-confidence SNP array-derived known sites and
- indexed for use in *gatk BaseRecalibrator*

**Table S1** Identification and distribution of 36,628 known SNP sites derived from the Illumina 60k *Brassica* SNP array across the 'Darmor-bzh' reference genome (v. 10)

| Chromosome | Length [bp] | Known Sites | |
|---|---|---|---|
| | | **SNP Count** | **bp between SNPs** |
| A01 | 32,958,928 | 1,763 | 18,695 |
| A02 | 33,432,960 | 1,444 | 23,153 |
| A03 | 39,685,748 | 2,290 | 17,330 |
| A04 | 23,101,715 | 1,541 | 14,991 |
| A05 | 42,112,164 | 1,727 | 24,385 |
| A06 | 45,146,386 | 1,527 | 29,565 |
| A07 | 29,390,523 | 2,043 | 14,386 |
| A08 | 26,309,499 | 1,361 | 19,331 |
| A09 | 53,549,826 | 1,848 | 28,977 |
| A10 | 20,778,245 | 1,692 | 12,280 |
| C01 | 48,239,358 | 2,703 | 17,847 |
| C02 | 62,297,340 | 2,934 | 21,233 |
| C03 | 73,669,886 | 2,888 | 25,509 |
| C04 | 65,837,619 | 3,528 | 18,661 |
| C05 | 56,382,805 | 1,110 | 50,795 |
| C06 | 50,218,839 | 1,505 | 33,368 |
| C07 | 55,656,957 | 2,158 | 25,791 |
| C08 | 41,681,856 | 1,347 | 30,944 |
| C09 | 66,465,249 | 1,219 | 54,524 |
| | | **36,628** | **23,438** |

**SNP array-derived variant sites improve base quality score recalibration in *B. napus***

1. Input data

- A recalibration file (csv) generated by *gatk AnalyzeCovariates*

2. Generation of bias plots before and after BQSR

Diagnostic plots for base quality scores, dinucleotide context, and machine cycle covariate bias are plotted with a custom R script:

- Plot_BQSR_Results_Minicore_Round1.r

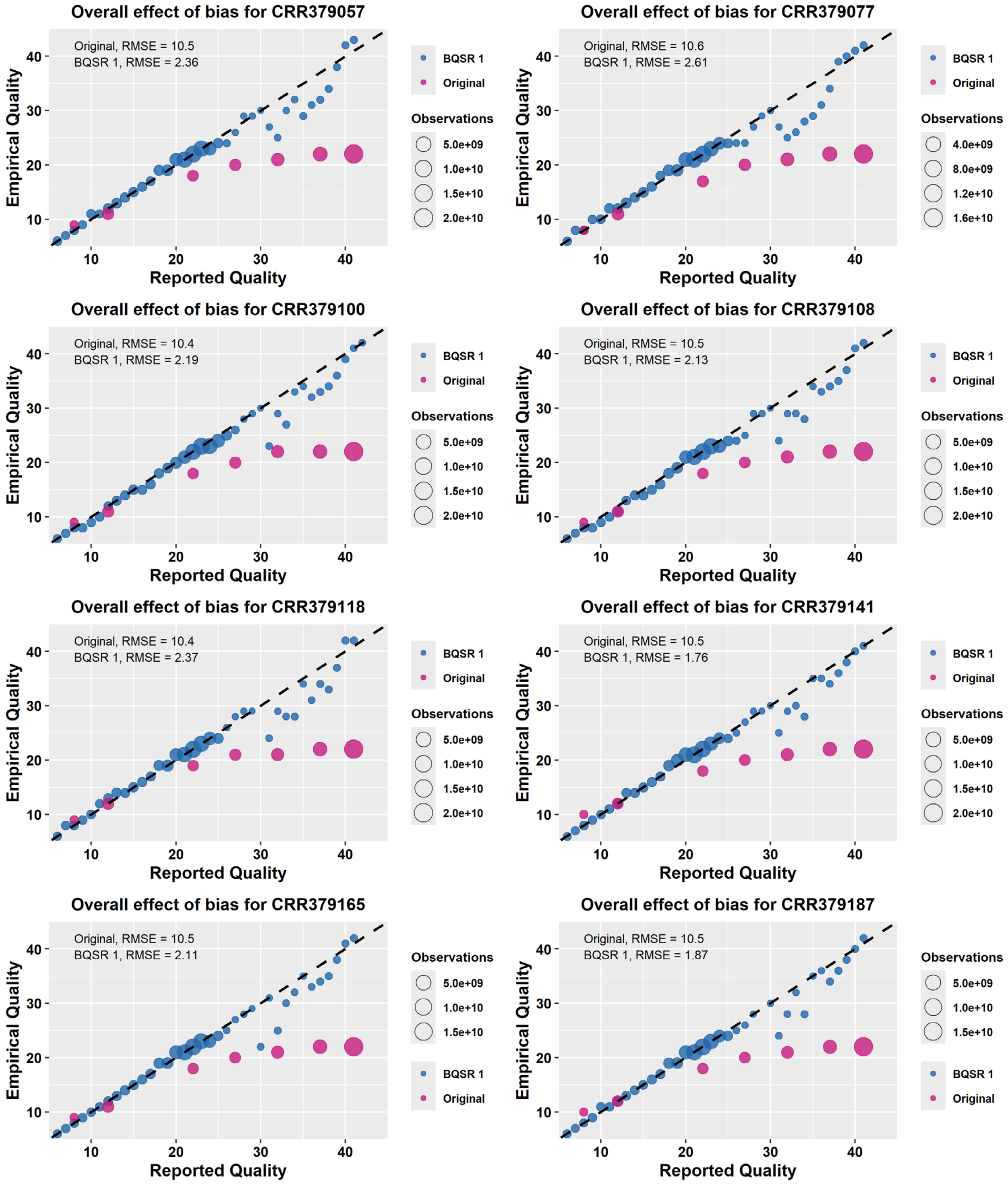


**Figure S1** Base quality score recalibration (BQSR) with the 36k known sites in the *Brassica napus* Damor-bzh reference genome (v. 10) for the base quality score bias across eight randomly selected samples from the *Brassica napus* mini-core collection. The raw uncalibrated data contains base quality scores in seven bins: Q8, Q12, Q22, Q27, Q32, Q37, Q41.

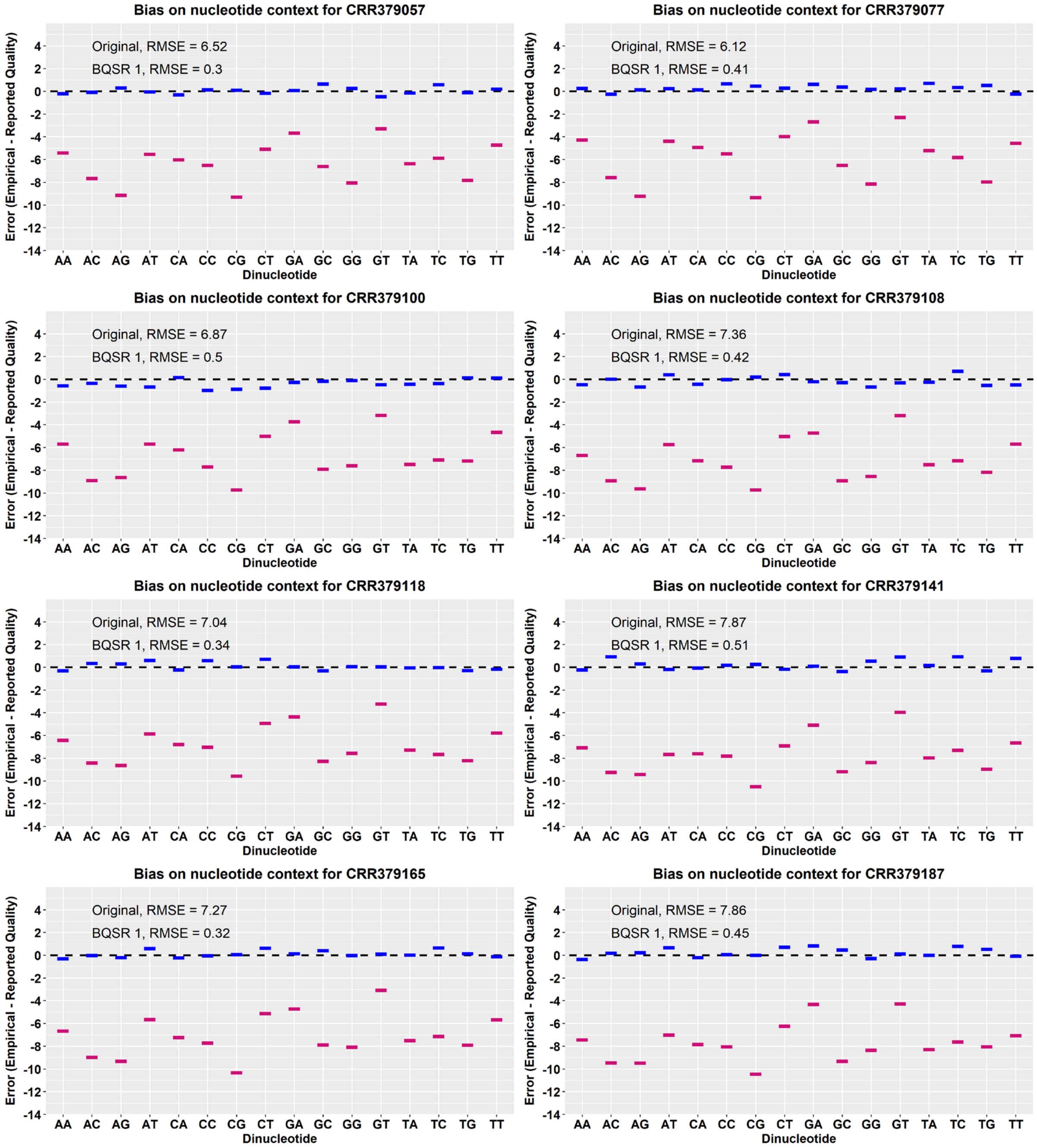


**Figure S2** Base quality score recalibration (BQSR) with the 36k known sites in the *Brassica napus* Darmor-bzh reference genome for the dinucleotide context bias across eight randomly selected samples from the *Brassica napus* mini-core collection. Pink = uncalibrated data; blue = recalibrated data.

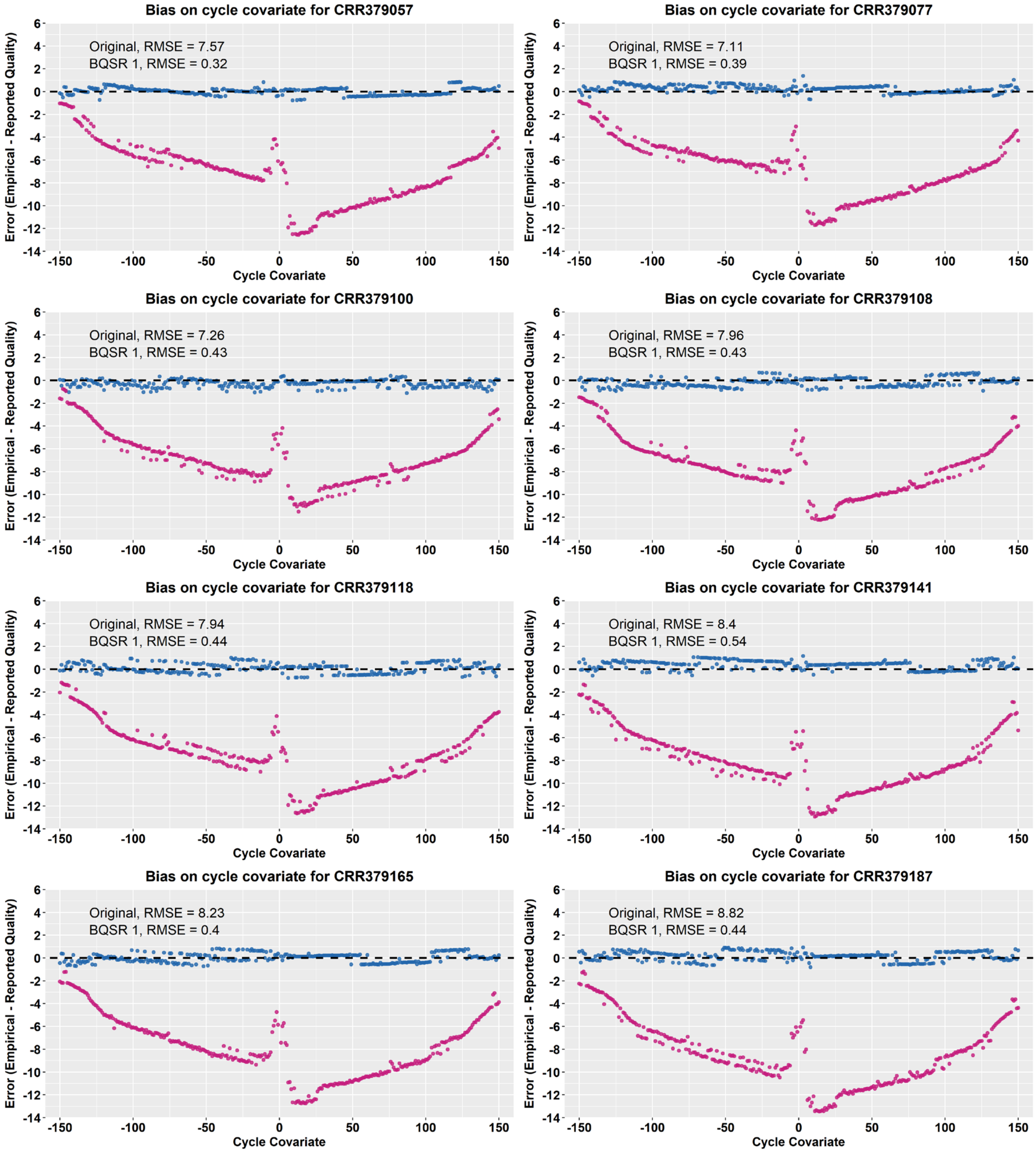


**Figure S3** Base quality score recalibration (BQSR) with the 36k known sites in the *Brassica napus* ‘Darmor-bzh’ reference genome for the machine cycle covariate bias across eight randomly selected samples from the *Brassica napus* mini-core collection. Pink = uncalibrated data; blue = recalibrated data.

#### Constructing a high-confidence truth set using SNP array-derived variant sites

After data pre-processing and the first round of variant calling (Table S2), matching variant sites between the unfiltered VCF and the 36k known sites are retrieved to create a cohort-specific truth set. This identifies 34,784 out of 25,107,582 SNP sites in the unfiltered VCF as true variant sites in the *Brassica napus* mini-core collection, hereafter referred to as 34k truth set. The exact commands to retrieve the truth set are provided below.

**Table S2** Variant calling summary in the *Brassica napus* mini-core collection

| Feature | Unfiltered VCF | 34k Truth Set |
|---|---|---|
| Total variants | 31,356,072 | 34,784 |
| SNP loci | 25,107,582 | 34,784 |
| INDEL loci | 5,601,848 | - |
| Ti/Tv-Ratio | 1.40 | 2.31 |

*Statistics were generated with gatk VariantEval*

1. Input data

- The unfiltered VCF of the cohort: *$RAW_SNP_VCF*
- The 36k known sites VCF: *$KNOWN_SITES_VCF*

2. Creating the 34k truth set with bcftools:

**###**

*# Extract positions from the 36k known sites VCF*

*$ bcftools query -f'%CHROM\t%POS\n' $KNOWN_SITES_VCF > $KNOWN_POS_TXT*

### Extract SNP sites from the unfiltered VCF that match the 36k known sites

*$ bcftools view – T $KNOWN_POS_TXT $RAW_SNP_VCF -Oz -o $TRUTH_SET_VCF*

### Index the Truth Set VCF

*$ bcftools index -t $TRUTH_SET_VCF*

**###**

Note: The SNP array-derived sites can also be extracted from the tab-delimited annotation table of the unfiltered VCF with *awk* as shown in the next section.

#### Preparing one-million-variant clustering subsets for annotation analysis

Subsets with one million randomly selected SNP and INDEL positions from the cohort VCF are generated for variant context annotation analysis and clustering.

Here, we demonstrate how to generate a SNP subset from the cohort VCF (Table S3), the same workflow applies for the INDEL subset by replacing 'SNP' with 'INDEL' below.

1. Input data

- The unfiltered VCF of the cohort: *$RAW_VCF*

2. Creating the clustering subset:

```
###
# Convert unfiltered VCF to tab-delimited annotation table:
$ gatk VariantsToTable -V $RAW_VCF   -F CHROM -F POS -F TYPE -F QD -F FS -F SOR -F MQ \
        -F MQRankSum -F ReadPosRankSum -O $RAW_TABLE
# Extract SNP positions from annotation table:
$ awk 'NR > 1 && $3 ~ /SNP/ {print $1 "\t" $2}' $RAW_TABLE > $SNP_POSITIONS
# Random shuffle of SNP positions and sorting afterwards:
$ shuf -n 1000000 $SNP_POSITIONS | sort -k1,1 -k2,2n > $SNP_POSITIONS_SUBSET
# Check distribution of SNP subset positions:
$ awk '{print $1}' $SNP_POSITION_SUBSET | sort | uniq -c
# Create the SNP subset:
$ awk 'FNR==NR{
        if(FNR>1) a[$1"\t"$2]
        next
}
FNR==1{print;next}
(($1"\t"$2) in a)' $SNP_POSITIONS_SUBSET $RAW_TABLE \
        > SNP_SUBSET_ANNOTATIONS
###
```

**Table S3** Summary of SNP distribution in the one-million-variant clustering subset of the *Brassica napus* mini-core collection

| Chromosome | Length [Mb] | Clustering Subset | |
|---|---|---|---|
| | | **SNP Count** | **bp between SNP** |
| A01 | 32,958,928 | 47,251 | 698 |
| A02 | 33,432,960 | 42,681 | 783 |
| A03 | 39,685,748 | 52,318 | 759 |
| A04 | 23,101,715 | 33,647 | 687 |
| A05 | 42,112,164 | 68,574 | 614 |
| A06 | 45,146,386 | 58,071 | 777 |
| A07 | 29,390,523 | 39,054 | 753 |
| A08 | 26,309,499 | 34,053 | 773 |
| A09 | 53,549,826 | 68,724 | 779 |
| A10 | 20,778,245 | 28,573 | 727 |
| C01 | 48,239,358 | 53,714 | 898 |
| C02 | 62,297,340 | 70,506 | 884 |
| C03 | 73,669,886 | 76,076 | 968 |
| C04 | 65,837,619 | 71,216 | 924 |
| C05 | 56,382,805 | 50,358 | 1,120 |
| C06 | 50,218,839 | 50,312 | 998 |
| C07 | 55,656,957 | 53,152 | 1,047 |
| C08 | 41,681,856 | 40,281 | 1,035 |
| C09 | 66,465,249 | 61,439 | 1,082 |
| | | **1,000,000** | **846** |

#### Comparing variant annotation profiles between call sets

The one-million-variant clustering subset and the 34k truth set are loaded into R and variants overlapping with the truth set are marked as true variants in the clustering subset. We plot the distribution of the variant context annotations for the clustering subset and for the 34k truth set (Figure S4). The same workflow applies for the INDEL subset without a truth set.

1. Input data

- The clustering subset of the unfiltered VCF of the cohort
- The 34k truth set derived from the unfiltered VCF of the cohort

2. Analysing variant context annotation profiles of the VCFs:

The variants are loaded into R and their variant context annotation profiles are analysed in part 1 of the R script:

- Variant_Context_Annotation_Analysis_Clustering_Round1_DV10_SNPs.r

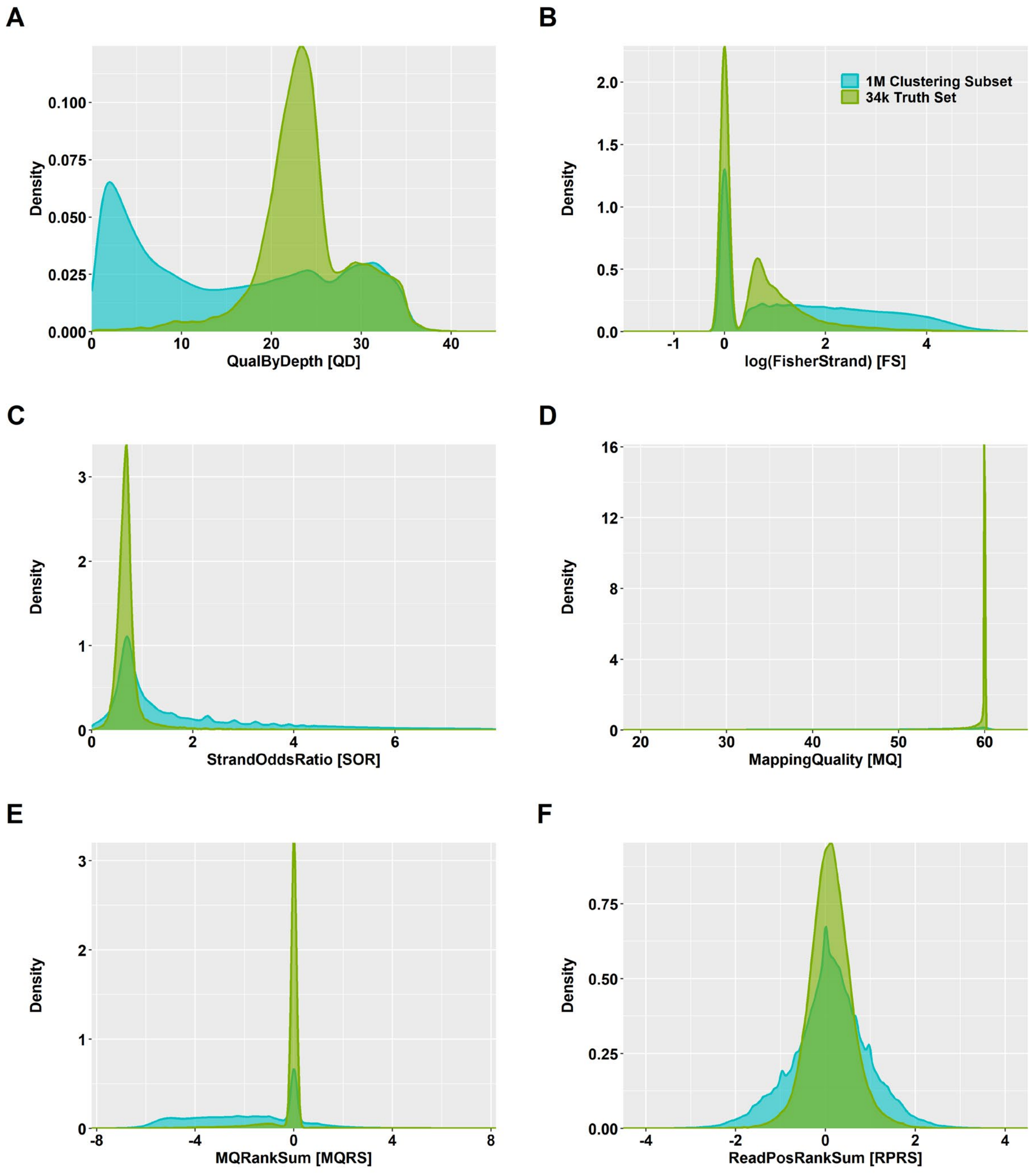


**Figure S4**.Density plots for **A** QualByDepth (QD), **B** FisherStrand (FS), **C** StrandOddsRatio (SOR), **D** MappingQuality (MQ), **E** MQRankSum (MQRS), and **F** ReadPosRankSum (RPRS) in the one-million-variant clustering subset (blue) and the 34k truth set (green) in the *Brassica napus* mini-core collection.

#### Characterization of the multi-dimensional annotation space for clustering

The annotations of the one-million-variant clustering subset are Z-scaled and the multi-dimensional relationships among the six annotations are analysed via principal component analysis (PCA) with a subset of 103,237 variants.

The same workflow applies for the INDEL subset.

1. Input data

- The clustering subset of the unfiltered VCF of the cohort

2. Multi-dimensional variant context annotation analysis via PCA:

The multi-dimensional relationships among the six annotations in the clustering subset are analysed in part 2 of the R script:

- Variant_Context_Annotation_Analysis_Clustering_Round1_DV10_SNPs.r

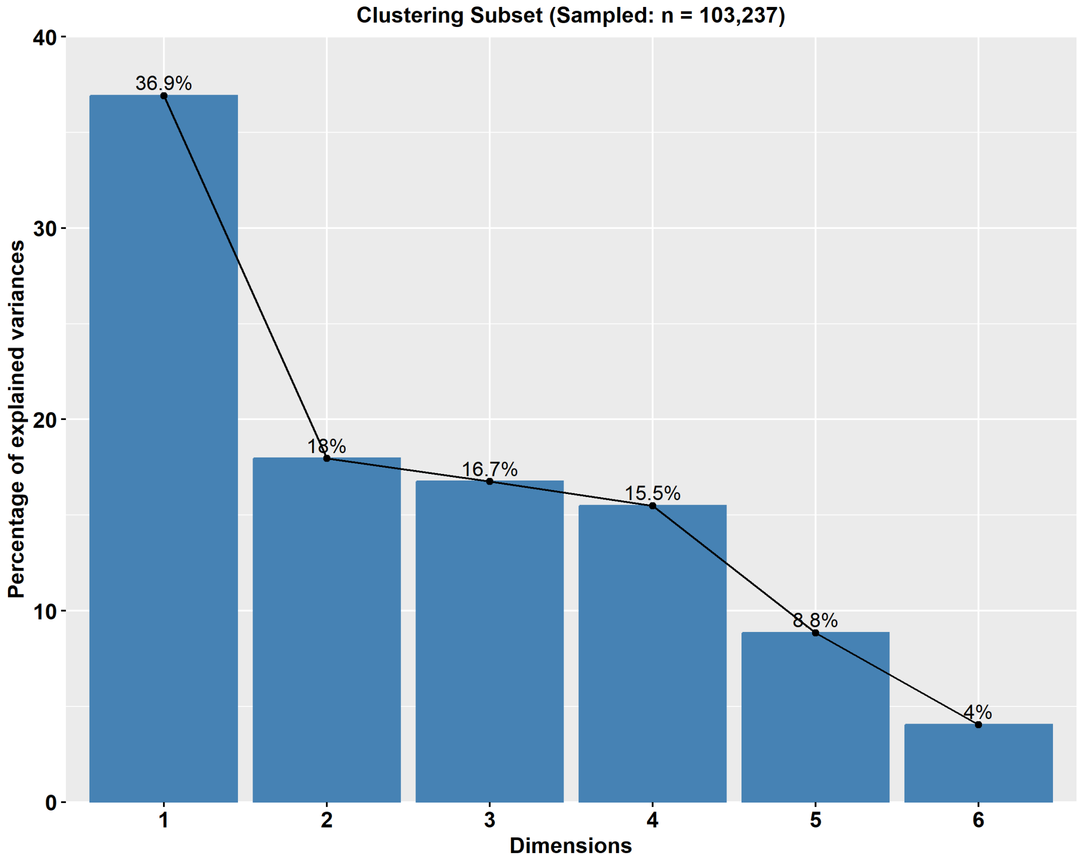


**Figure S5** Scree plots of principal components for six variant context annotations in the one-million-variant clustering subset from the *Brassica napus* mini-core collection. The principal component analysis was performed with complete observations for QualByDepth, Fisher Strand, Strand Odds Ratio, Mapping Quality, MQRankSum, and ReadPosRankSum.

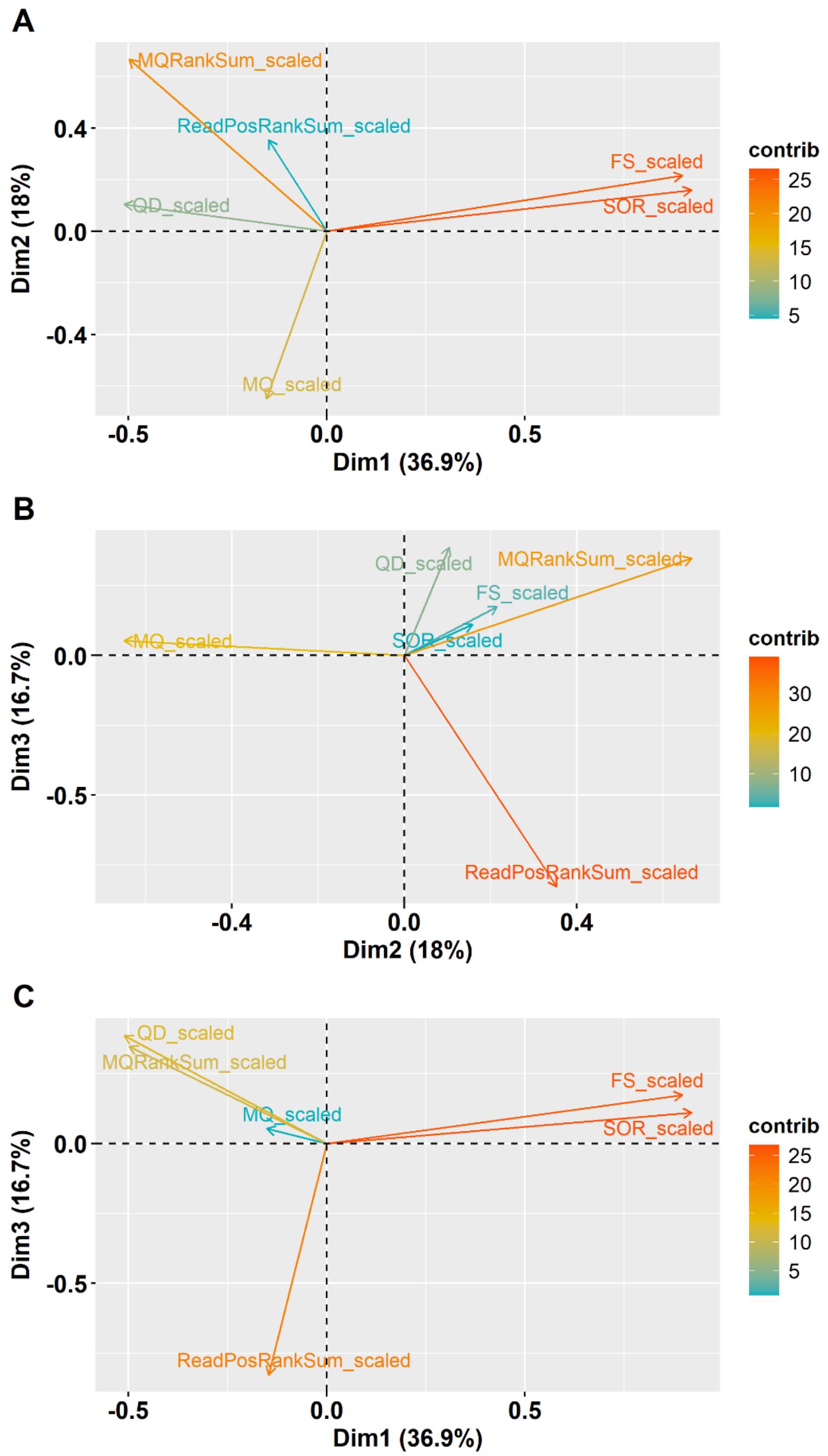


**Figure S6**.Loading plots of the principal components (PCs) based on ~100k SNPs from the one-million-variant clustering subset of the *Brassica napus* mini-core collection. **A – C** Loading plots for PCs 1 – 3 with six variant context annotations. *QD* QualByDepth *FS* FisherStrand *SOR* StrandOddsRatio *MQ* MappingQuality

##### Unsupervised GMM clustering identifies high-quality variant classes

We fit a Gaussian mixture model (GMM) with three components based on the ΔBIC method.

1. Input data

- The clustering subset of the unfiltered VCF of the cohort

2. GMM-based variant context annotation clustering:

The GMM-based clustering is performed in part 3 of the R script:

- Variant_Context_Annotation_Analysis_Clustering_Round1_DV10_SNPs.r

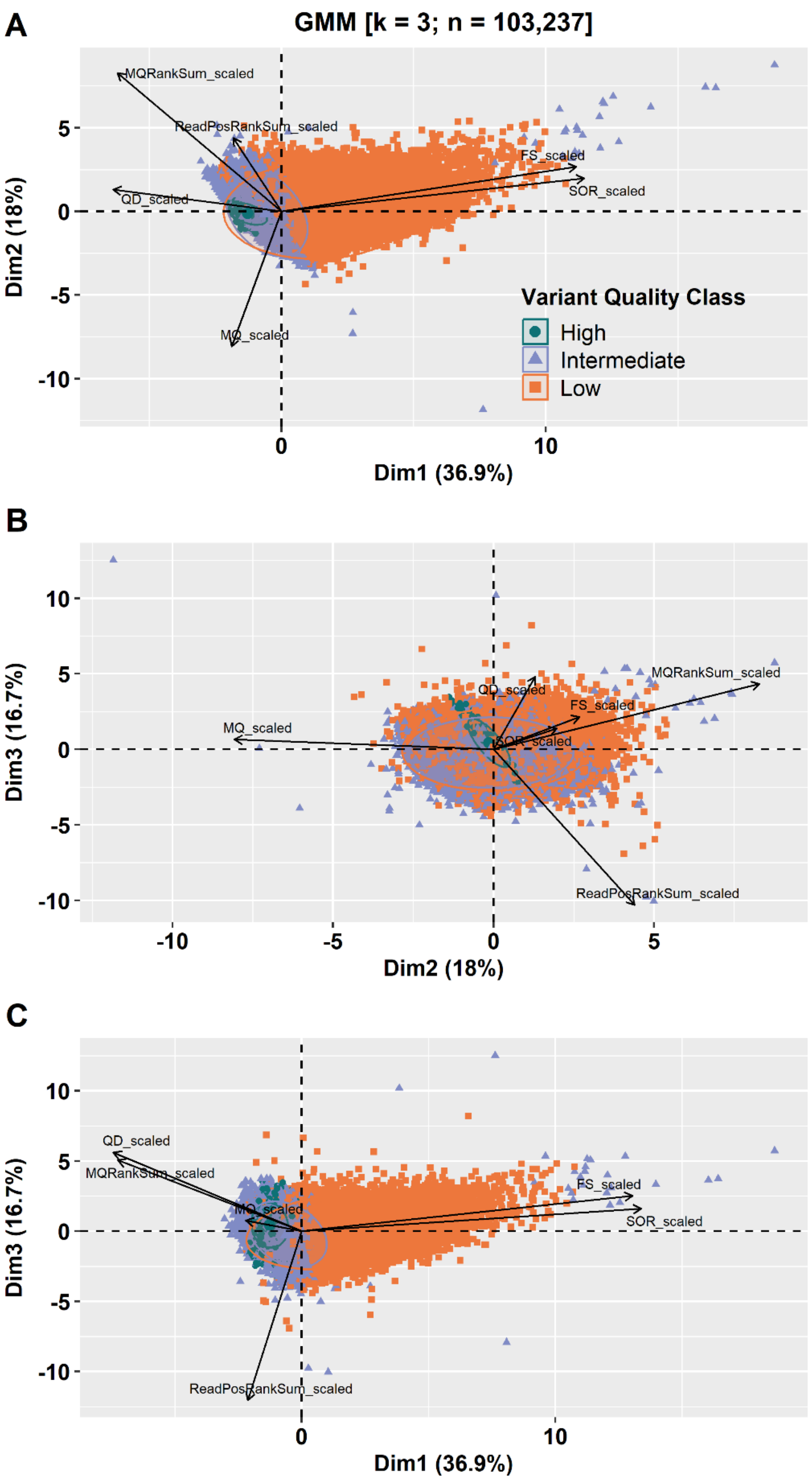


**Figure S7** Principal component analysis (PCA) biplots for 103,237 variants from the one-million-variant clustering subset of the *Brassica napus* mini-core collection. Variant annotation variables are shown as loading vectors. Panels **A – C** show variants coloured according to variant quality class based on the three-component Gaussian mixture model clustering. Green = high, purple = intermediate, orange = low.

#### Construction of cohort-specific truth and training resources for VQSR

1. Input data

- The clustering subset of the unfiltered VCF of the cohort

2. Training set construction:

Variant stratification by posterior probability within each cluster and annotation profile analysis is performed in part 4 of the R script:

- Variant_Context_Annotation_Analysis_Clustering_Round1_DV10_SNPs.r

**Table S4** Summary statistics for SNP stratification by posterior probability across the three variant quality clusters in the *Brassica napus* mini-core collection

***High-Quality Cluster***

| Posterior Probability [%] | n | Med(x) | | | | | | |
|---|---|---|---|---|---|---|---|---|
| | | QD | FS | SOR | MQ | MQRS | RPRS | HFV |
| 0-25 | 0 | - | - | - | - | - | - | - |
| 25-50 | 48 | 9.5 | 8 | 1.9 | 60 | 0 | 0.2 | 7 |
| 50-60 | 171 | 19.4 | 4.7 | 1.0 | 59 | 0 | 0.1 | 11 |
| 60-70 | 190 | 19.9 | 4.0 | 0.9 | 59 | 0 | 0.0 | 4 |
| 70-80 | 220 | 18.9 | 4.9 | 0.9 | 59 | 0 | 0.0 | 11 |
| 80-90 | 405 | 20.4 | 5.3 | 1.0 | 59 | 0 | 0.1 | 19 |
| 90-100 | 146,777 | 22.6 | 0.9 | 0.7 | 60 | 0 | 0.1 | 1,086 |

***Intermediate-Quality Cluster***

| Posterior Probability [%] | n | Med(x) | | | | | | |
|---|---|---|---|---|---|---|---|---|
| | | QD | FS | SOR | MQ | MQRS | RPRS | HFV |
| 0-25 | 0 | - | - | - | - | - | - | - |
| 25-50 | 340 | 11.8 | 49.1 | 3.4 | 40 | 0 | 0.2 | 592 |
| 50-60 | 4,827 | 7.0 | 10.0 | 1.5 | 53 | -2.6 | 0.1 | 1,627 |
| 60-70 | 5,544 | 7.7 | 9.8 | 1.5 | 53 | -2.5 | 0.1 | 1,780 |
| 70-80 | 7,320 | 7.7 | 9.4 | 1.4 | 53 | -2.6 | 0.1 | 2,379 |
| 80-90 | 13,330 | 7.2 | 8.6 | 1.4 | 53 | -2.7 | 0.1 | 4,283 |
| 90-100 | 383,199 | 14.2 | 1.7 | 0.7 | 54 | -1.9 | 0.2 | 101,417 |

***Low-Quality Cluster***

| Posterior Probability [%] | n | Med(x) | | | | | | |
|---|---|---|---|---|---|---|---|---|
| | | QD | FS | SOR | MQ | MQRS | RPRS | HFV |
| 0-25 | 0 | - | - | - | - | - | - | - |
| 25-50 | 4,544 | 15.1 | 190.0 | 8.8 | 43 | -2.1 | 0.0 | 10,104 |
| 50-60 | 5,188 | 7.4 | 11.1 | 1.6 | 52 | -2.5 | 0.1 | 2,984 |
| 60-70 | 5,423 | 7.8 | 11.4 | 1.7 | 52 | -2.5 | 0.1 | 3,145 |
| 70-80 | 6,403 | 8.0 | 11.8 | 1.7 | 52 | -2.4 | 0.1 | 4,000 |
| 80-90 | 9,478 | 8.4 | 12.3 | 1.8 | 52 | -2.2 | 0.1 | 6,480 |
| 90-100 | 304,614 | 5.4 | 27.8 | 3.2 | 53 | -3.0 | 0.0 | 321,246 |

*n* number of variants; *QD* QualByDepth; *FS* FisherStrand; *SOR* StrandOddsRatio; *MQ* MappingQuality; *MQRS* MappingQualityRankSum; *RPRS* ReadPosRankSum; *HFV* Hard filter violation rate

**Table S5** Summary statistics for INDEL quality classes based on GMM clustering in the *Brassica napus* mini-core collection

| INDEL Set | Variant Class | #INDELs | *Med(x)* | | | | |
|---|---|---|---|---|---|---|---|
| | | | QD | FS | SOR | MQRS | RPRS |
| Clustering Subset (*k* = *4*) | Very High | 220,351 | 29.9 | 0.9 | 0.7 | 0 | 0.1 |
| | High | 200,377 | 9.4 | 1.1 | 0.7 | -1 | 0 |
| | Intermediate | 284,326 | 11.4 | 9.5 | 1.6 | -2 | 0 |
| | Low | 163,188 | 6.4 | 45.9 | 4.7 | -2.9 | -0.2 |

*k* number of clusters; *QD* QualByDepth; *FS* FisherStrand; *SOR* StrandOddsRatio; *MQRS* MappingQualityRankSum; *RPRS* ReadPosRankSum

**Table S6** Summary statistics for cohort-specific INDEL quality score recalibration across different truth sensitivity tranches in the *Brassica napus* mini-core collection

| Truth Sensitivity Tranche | Known Sites | Novel Sites | Known Sites [Ti/Tv] | Novel Sites [Ti/Tv] | minVQSLod | Model | Accessible Truth Sites | Calls Truth Sites |
|---|---|---|---|---|---|---|---|---|
| 90.00 | - | 1,562,649 | - | - | 3.59 | INDEL | 388,346 | 349,511 |
| 95.00 | - | 1,865,619 | - | - | 3.20 | INDEL | 388,346 | 368,928 |
| 99.00 | - | 2,737,955 | - | - | 2.22 | INDEL | 388,346 | 384,462 |
| 99.50 | - | 3,098,067 | - | - | 1.89 | INDEL | 388,346 | 386,404 |
| 99.70 | - | 3,373,845 | - | - | 1.58 | INDEL | 388,346 | 387,180 |
| 99.90 | - | 3,721,637 | - | - | 1.10 | INDEL | 388,346 | 387,957 |
| 99.99 | - | 4,095,553 | - | - | 0.62 | INDEL | 388,346 | 388,307 |
| 100.00 | - | 6,248,493 | - | - | -39832.96 | INDEL | 388,346 | 388,346 |

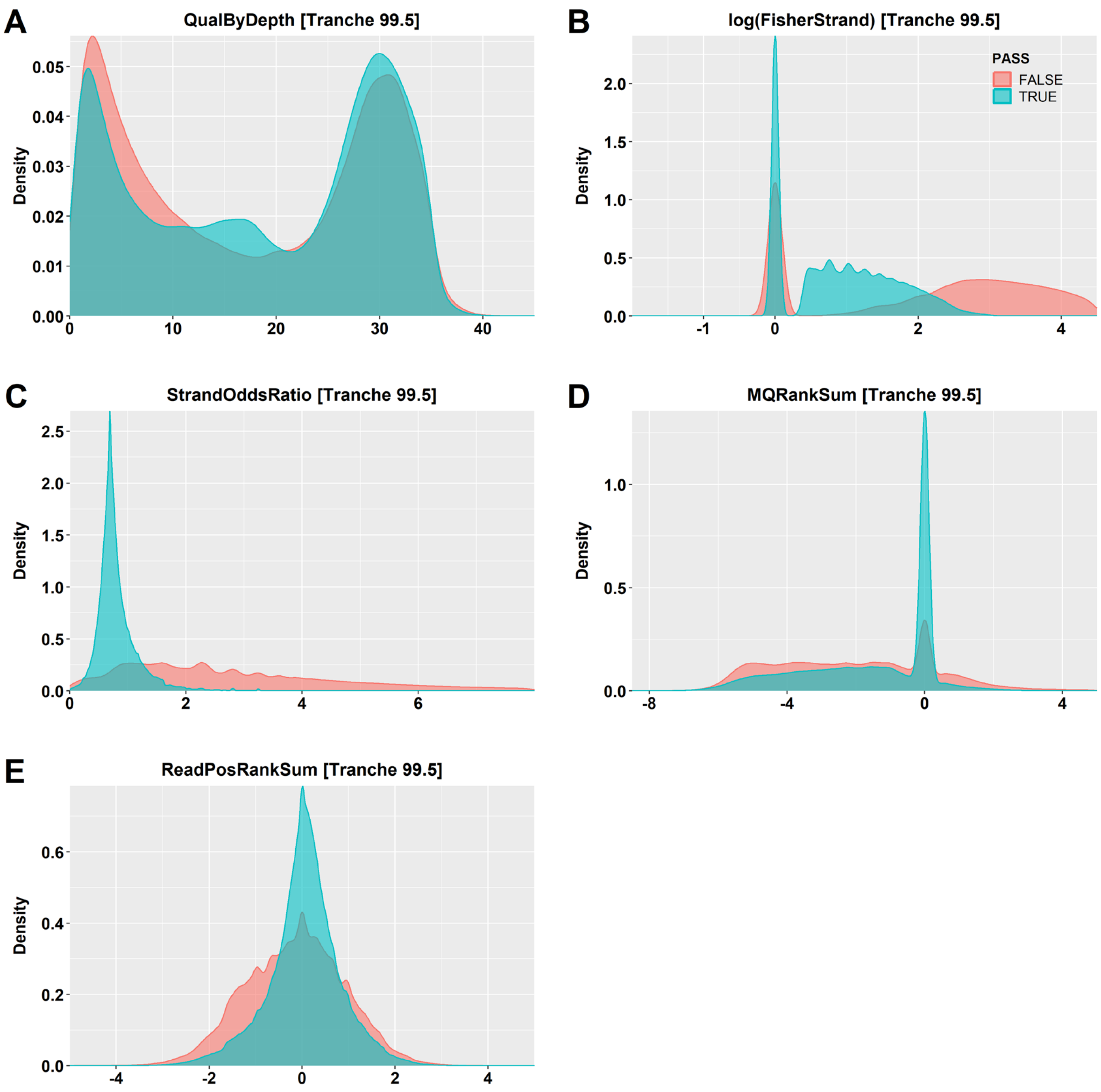


**Figure S8** Distribution of **A** QualByDepth, **B** log-transformed Fisher Strand, **C** Strand Odds Ratio, **D** MQRankSum, and **E** ReadPosRankSum values for retained (blue) and filtered (red) INDELs following variant quality score recalibration (VQSR) and filtering at the 99.5% tranche in a one-million-variant subset from the *Brassica napus* mini-core collection.
