## Supplementary Data 2 for "Unsupervised Clustering Enables Cohort-Specific Variant Recalibration in Crops"

### Unsupervised Clustering Enables Cohort-Specific Variant Recalibration in Crops

Thomas Bergmann[1,2], Tessa MacNish[1], Jacqueline Batley[2], and David Edwards[1]

[1]Centre for Applied Bioinformatics, University of Western Australia, 6009 Perth, Australia

[2]School of Biological Sciences, University of Western Australia, 6009 Perth, Australia

#### Variant Calling and Filtering in a Wild *Brassica oleracea* Population

<u>Scripts and resources:</u>

https://github.com/tbergmann92/ Advanced-Crop-Variant-Discovery

#### Preparation of SNP array-derived known variant sites in *Brassica oleracea*

The SNP array-derived known sites are prepared as described in Supplementary Data 1 with the corresponding *Brassica oleracea* ‘W03’ reference genome and the 4_Prepare_60k_SNP_Array_W03.sh script.

**Table S1** Identification and distribution of 26,317 known SNP sites derived from the Illumina 60k *Brassica* SNP array across the *Brassica oleracea* ‘W03’ reference genome

| Chromosome | Length [bp] | Known Sites | |
|---|---|---|---|
| | | SNP Count | bp between SNPs |
| C01 | 58,570,197 | 3,313 | 17,679 |
| C02 | 72,559,525 | 3,384 | 21,442 |
| C03 | 80,735,250 | 3,929 | 20,549 |
| C04 | 71,180,230 | 4,494 | 15,839 |
| C05 | 63,147,374 | 1,764 | 35,798 |
| C06 | 49,313,063 | 2,194 | 22,476 |
| C07 | 64,564,754 | 2,877 | 22,442 |
| C08 | 57,383,795 | 2,318 | 24,756 |
| C09 | 76,892,030 | 2,044 | 37,618 |
| | | **26,317** | **23,364** |

#### SNP array-derived variant sites improve base quality score recalibration in *B. oleracea*

1. Input data

- A recalibration file (csv) generated by *gatk AnalyzeCovariates*

2. Generation of bias plots before and after BQSR

Diagnostic plots for base quality scores, dinucleotide context, and machine cycle covariate bias are plotted with a custom R script:

- Plot_BQSR_Results_Wild_Bol.r

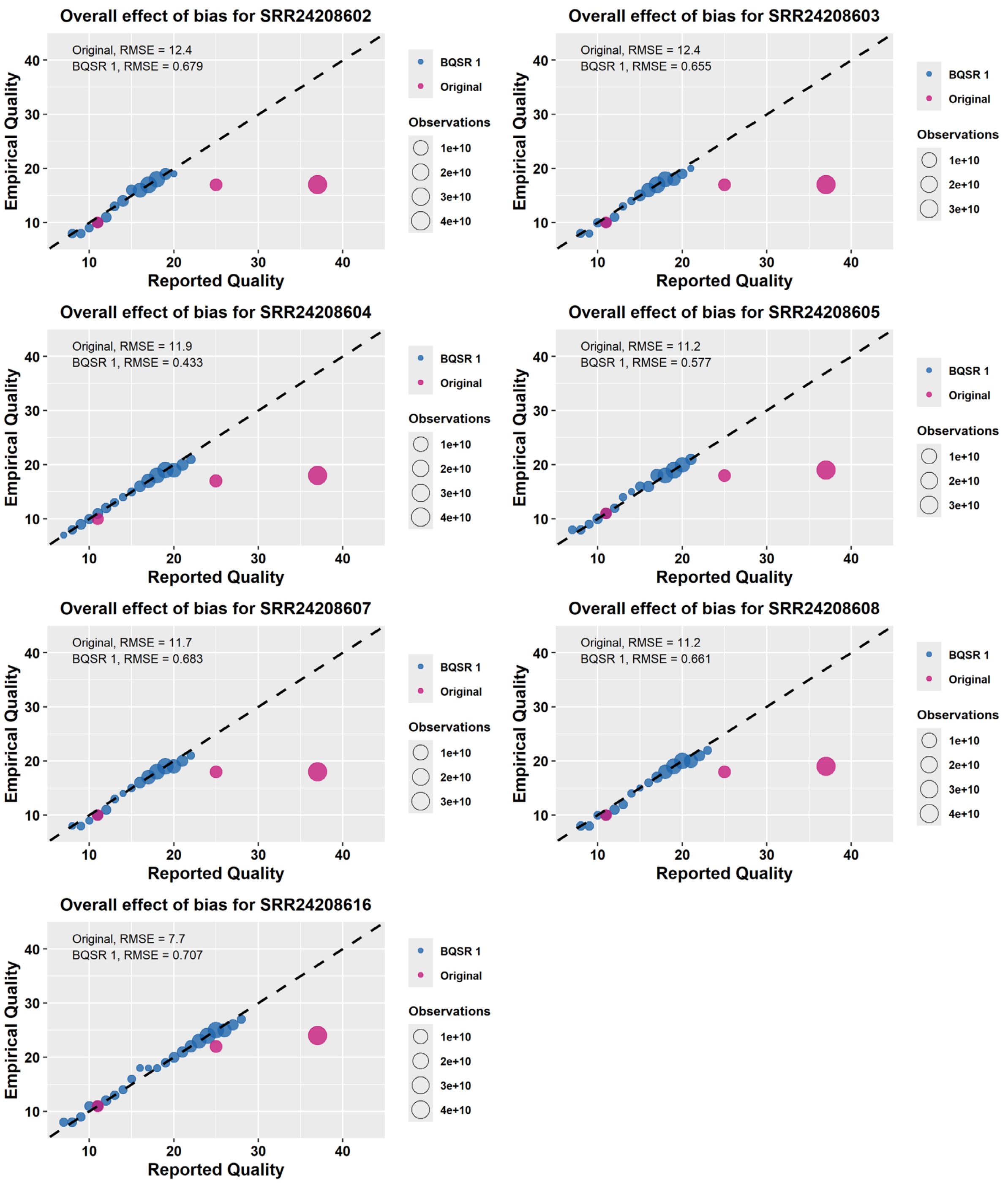


**Figure S1**.Base quality score recalibration (BQSR) with the 26k known sites in the *Brassica oleracea* 'W03' reference genome for the base quality score bias across seven samples from the wild *Brassica oleracea* collection. The raw uncalibrated data contains base quality scores in three bins: Q11, Q25, Q37. Pink = uncalibrated data; blue = recalibrated data.

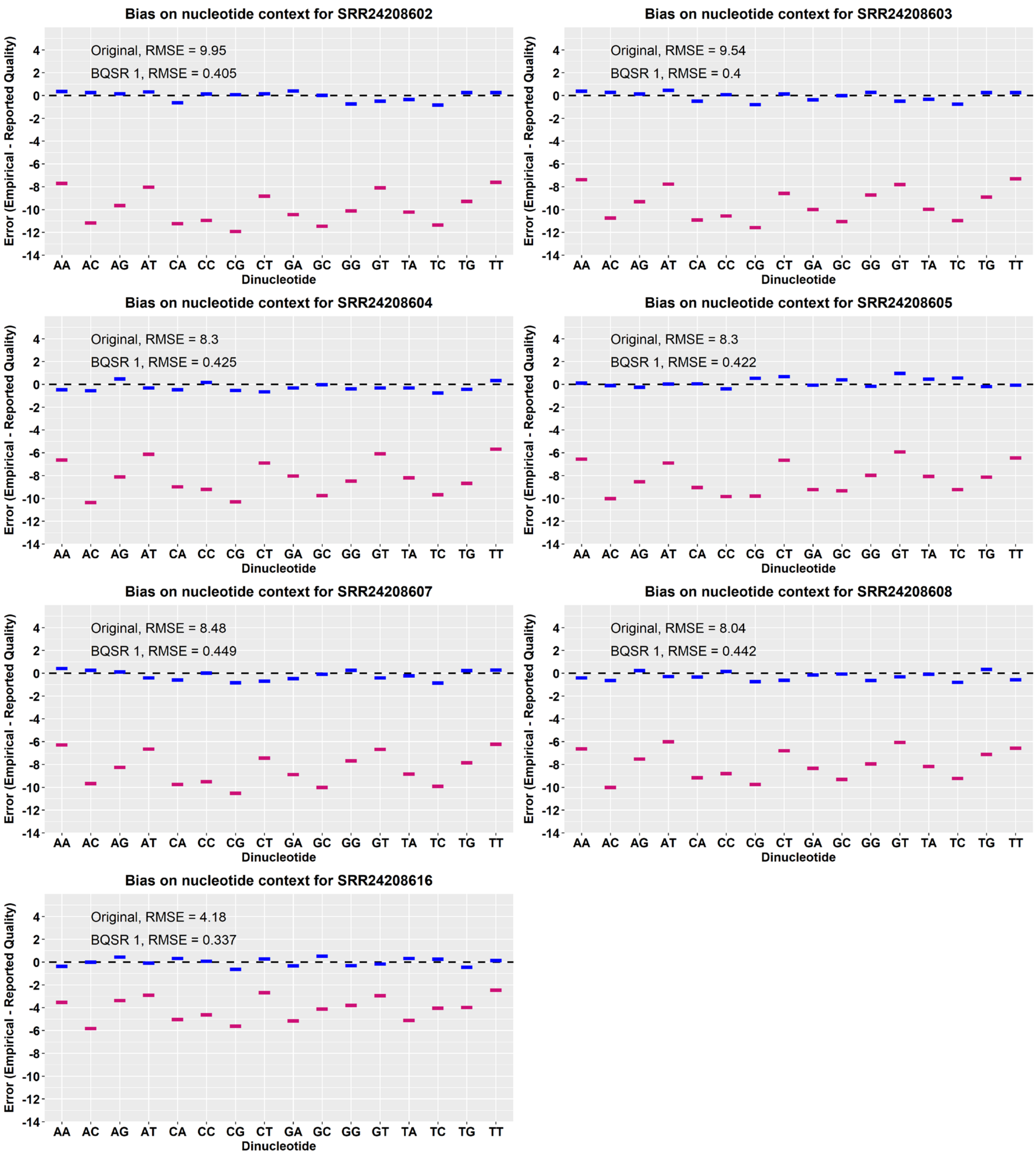


**Figure S2**.Base quality score recalibration (BQSR) with the 26k known sites in the *Brassica oleracea* ‘W03’ reference genome for the dinucleotide context bias across the seven samples from the wild *Brassica oleracea* collection. Pink = uncalibrated data; blue = recalibrated data.

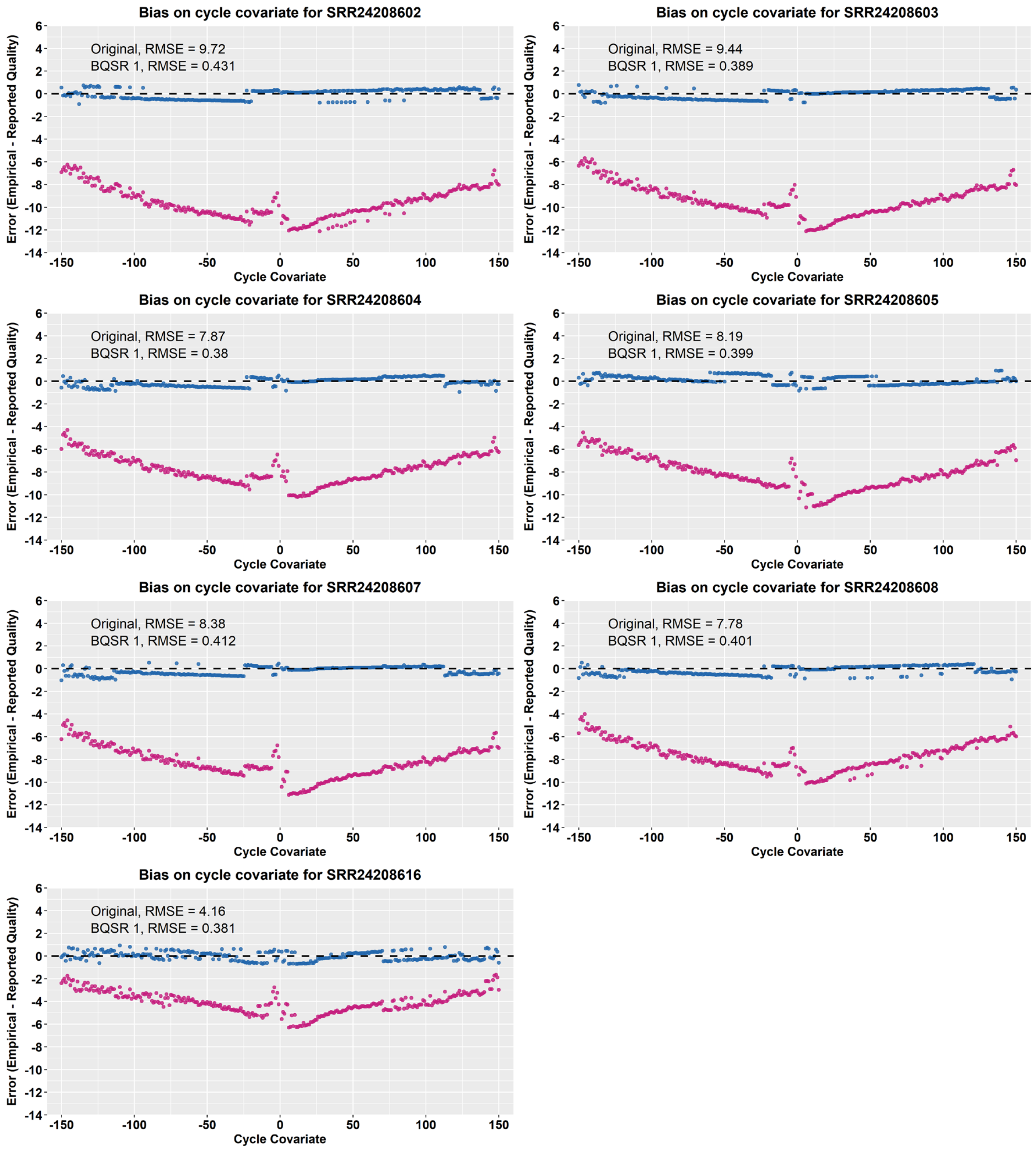


**Figure S3**.Base quality score recalibration (BQSR) with the 26k known sites in the *Brassica oleracea* ‘W03’ reference genome for the machine cycle covariate bias across seven samples from the wild *Brassica oleracea* collection. Pink = uncalibrated data; blue = recalibrated data.

#### Constructing a high-confidence truth set using SNP array-derived variant sites

After data pre-processing and the first round of raw variant calling (Table 2), matching variant sites between the unfiltered VCF and the 26k known sites are retrieved to create a cohort-specific truth set. This identifies 13,496 out of 23,828,683 SNP sites in the unfiltered VCF as true variant sites in the wild *Brassica oleracea* collection, hereafter referred to as 13k truth set.

**Table S2**.Variant calling summary in the wild *Brassica oleracea*.collection

| Feature | Raw VCF | 13k Truth Set |
|---|---|---|
| Total variants | 28,508,574 | 13,496 |
| SNP sites | 23,828,683 | 13,496 |
| INDEL sites | 4,220,052 | - |
| Ti/Tv-Ratio | 1.52 | 2.29 |

*Statistics were generated with gatk VariantEval*

> *Details on how to generate the truth set are found in Supplementary Data 1 in the corresponding section.*

#### Preparing one-million-variant clustering subsets for annotation analysis

> *Details on how to generate the clustering subset are found in Supplementary Data 1 in the corresponding section.*

**Table S3** Summary of SNP distribution in the one-million-variant clustering subset of the wild *Brassica oleracea* collection

| Chromosome | Length [Mb] | Clustering Subset | |
|---|---|---|---|
| | | SNP Count | bp per SNP |
| C01 | 58,570,197 | 100,232 | 584 |
| C02 | 72,559,525 | 124,948 | 581 |
| C03 | 80,735,250 | 135,570 | 596 |
| C04 | 71,180,230 | 116,881 | 609 |
| C05 | 63,147,374 | 103,455 | 610 |
| C06 | 49,313,063 | 84,564 | 583 |
| C07 | 64,564,754 | 106,694 | 605 |
| C08 | 57,383,795 | 96,730 | 593 |
| C09 | 76,892,030 | 130,926 | 587 |
| | | **1,000,000** | **857** |

#### Comparing variant annotation profiles between call sets

> *Details on how to analyse the variant context annotations are found in Supplementary Data 1 in the corresponding section.*

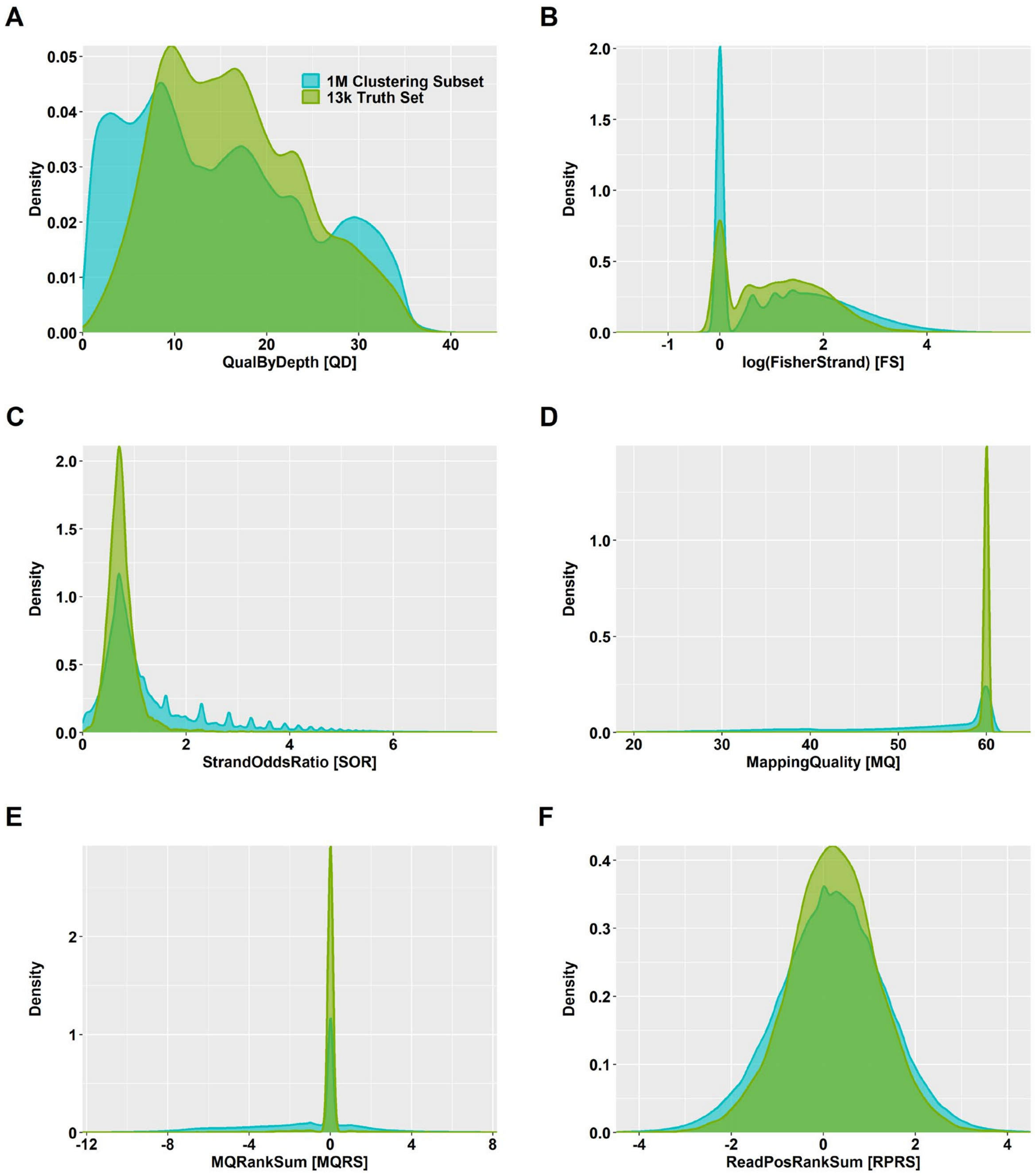


**Figure S4**.Density plots for **A** QualByDepth (QD), **B** Fisher Strand (FS), **C** Strand Odds Ratio (SOR), **D** Mapping Quality (MQ), **E** MQRankSum (MQRS), and **F** ReadPosRankSum (RPRS) in the one-million-variant clustering subset (blue) and the 13k truth set (green) in the wild *Brassica oleracea* population.

#### Characterization of the multi-dimensional annotation space for clustering

*> Details on how to analyse the multi-dimensional annotation space are found in Supplementary Data 1 in the corresponding section.*

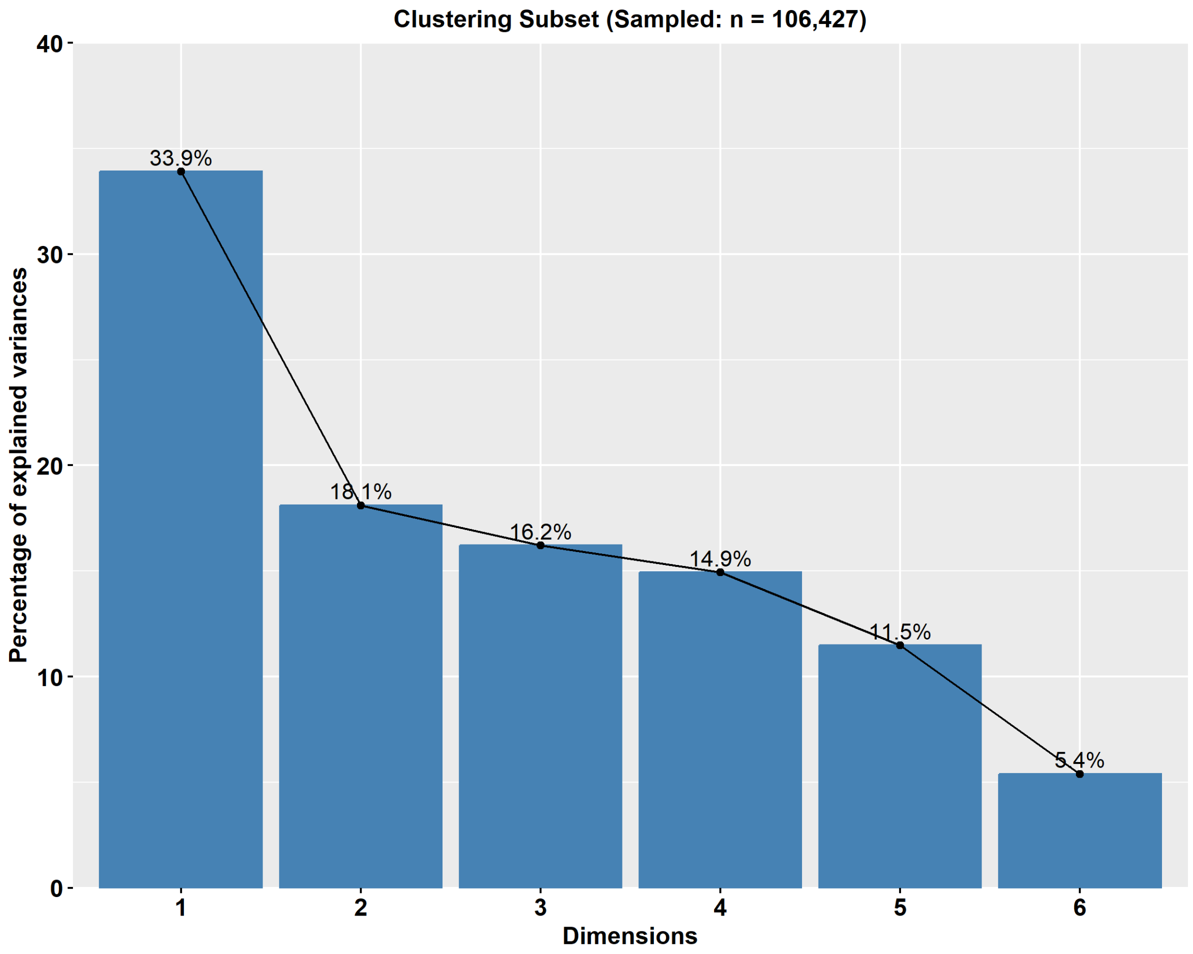


**Figure S5**.Scree plots of principal components for six variant context annotations in the one-million-variant clustering subset from the wild *Brassica oleracea* population. The principal component analysis was performed with complete observations for QualByDepth, Fisher Strand, Strand Odds Ratio, Mapping Quality, MQRankSum, and ReadPosRankSum.

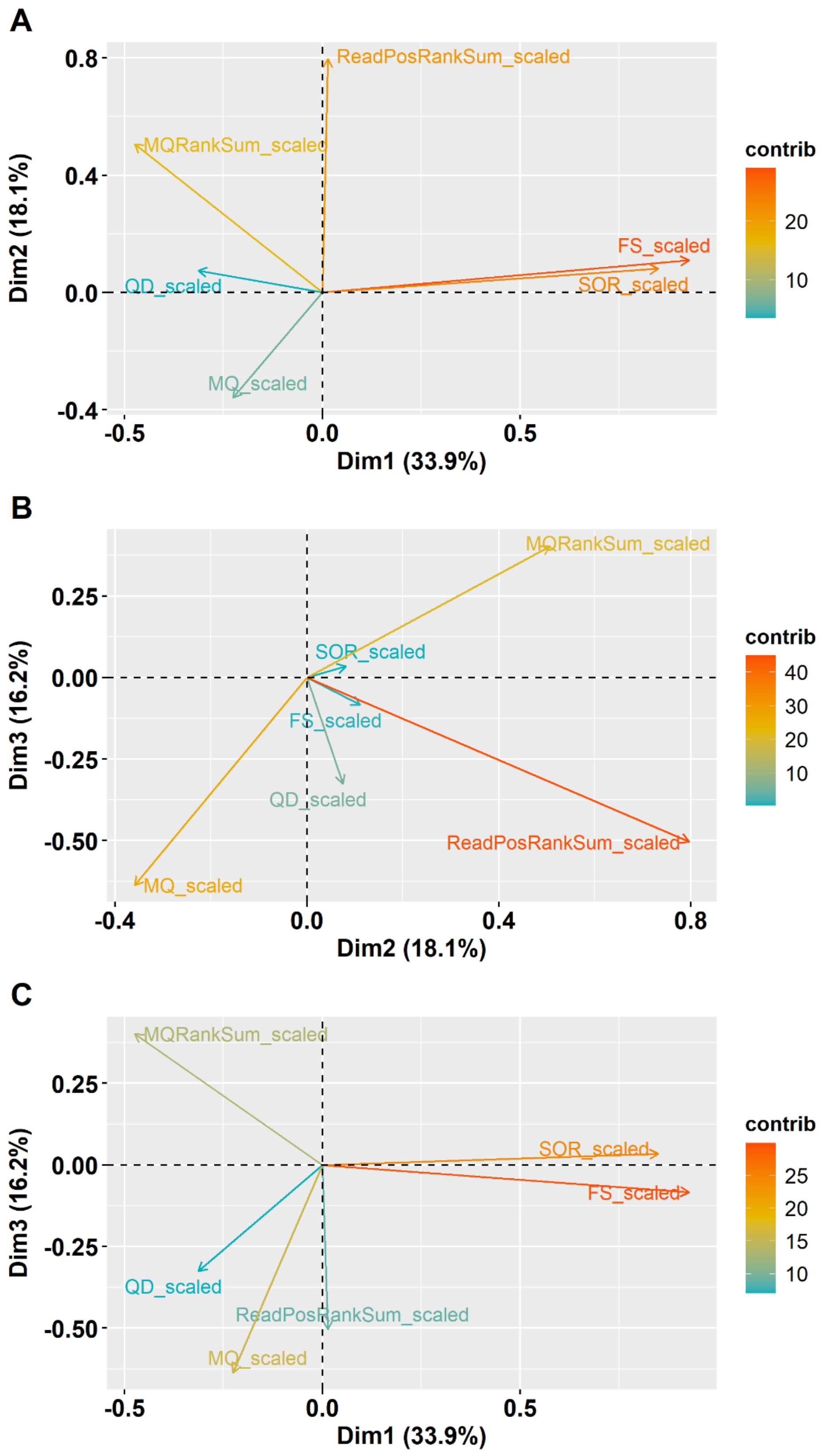


**Figure S6**.Loading plots of the principal components (PCs) based on ~100k SNPs from the one-million-variant clustering subset of the *Brassica oleracea* population. **A – C** Loading plots for PCs 1 – 3 with six variant context annotations. *QD* QualByDepth *FS* Fisher Strand *SOR* Strand Odds Ratio *MQ* Mapping Quality

#### Unsupervised GMM clustering identifies high-quality variant classes

*> Details on how to build the model are found in Supplementary Data 1 in the corresponding section.*

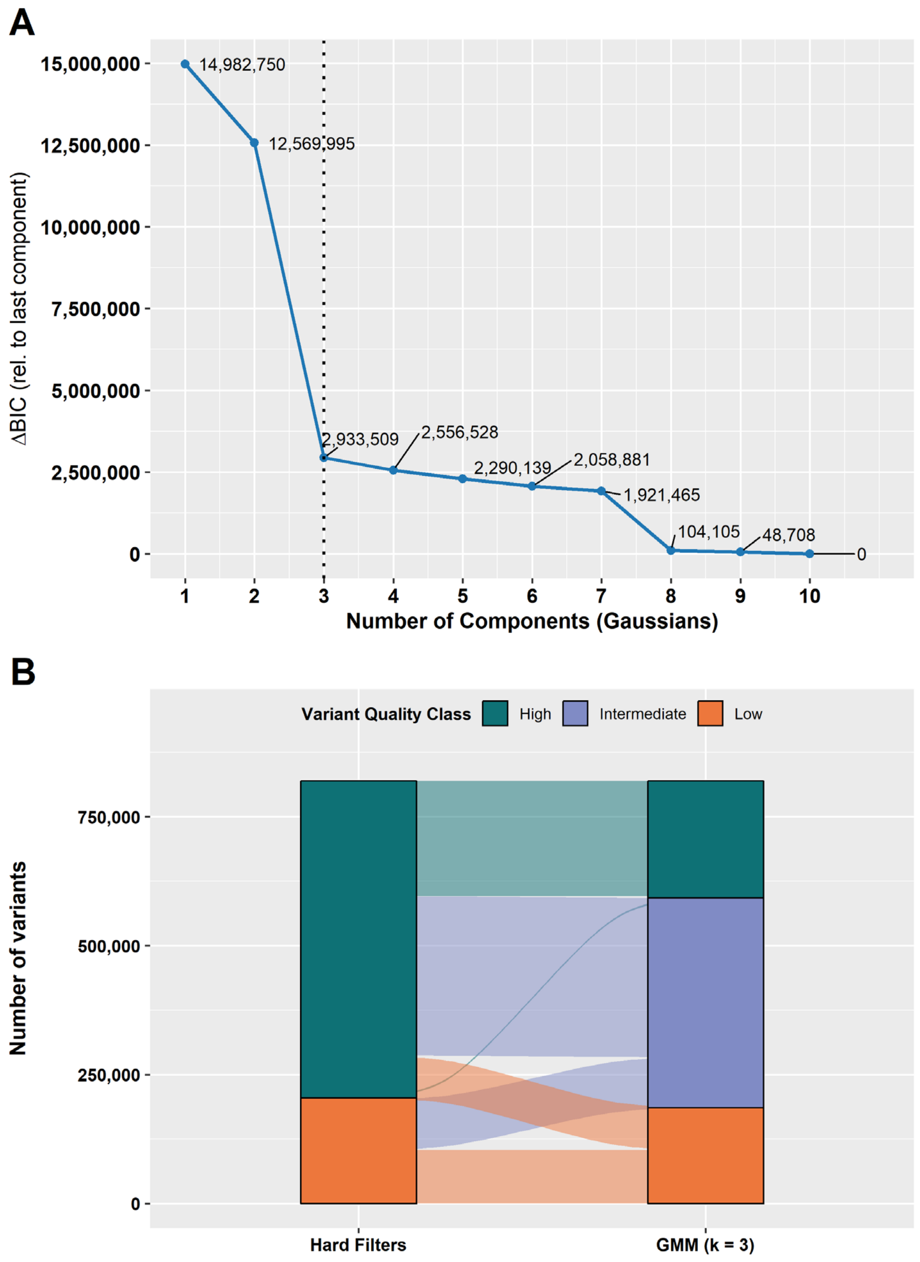


**Figure S7**.Model selection and variant quality class stratification using Gaussian mixture model (GMM)-based clustering. **A** Change in Bayesian Information Criterion (ΔBIC) relative to the largest model ($k$ = 10) across increasing numbers of Gaussian components ($k$). Lower values indicate better model support. The dashed vertical line indicates best-model-fit. **B** Alluvial plot showing the flow of variant classification between hard filtering and GMM-based clustering ($k$ = 3). Each stratum represents the classification at a given stage, and its vertical width corresponds to the number of variants. Alluvia are coloured according to group assignments from the GMM with $k$ = 3.

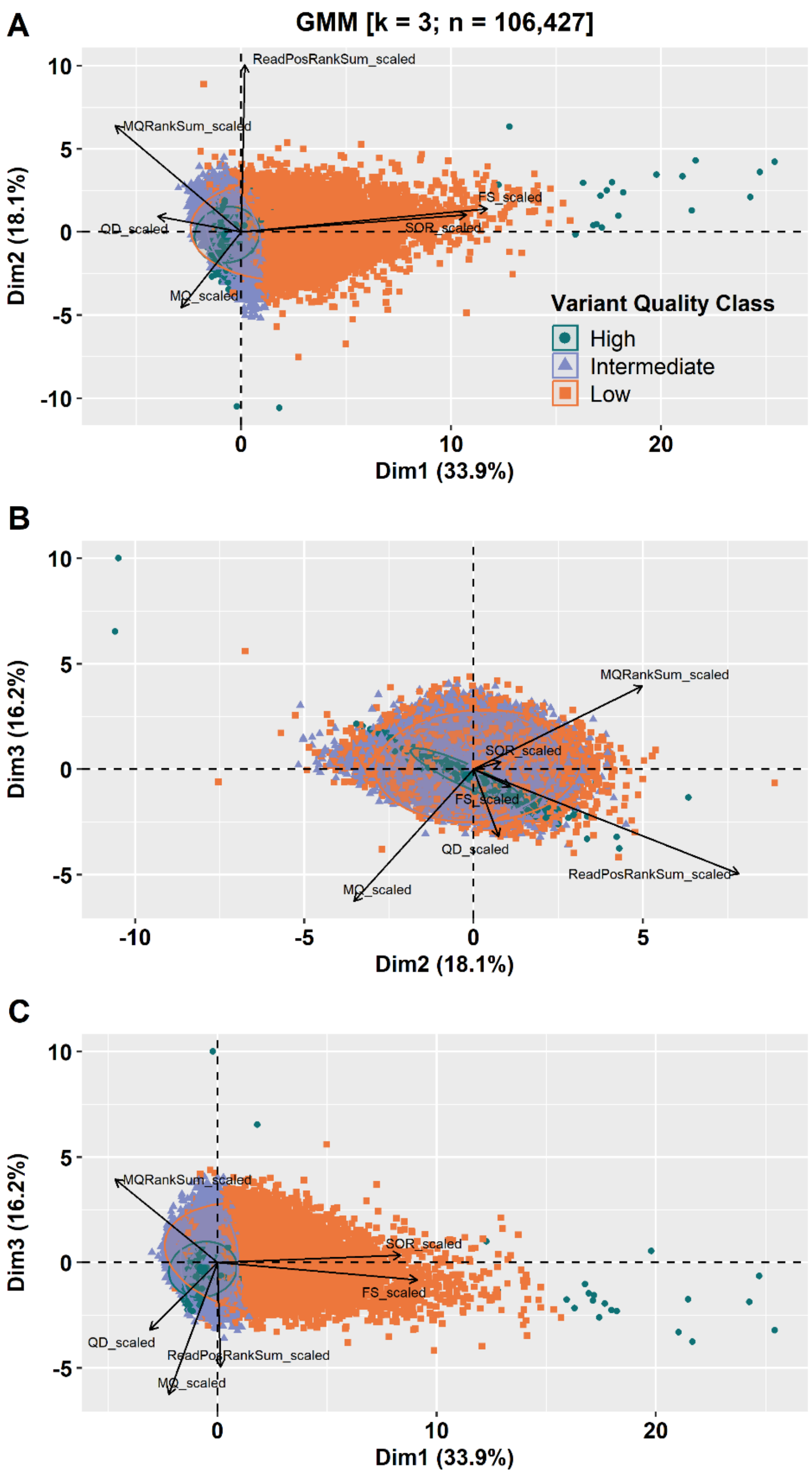


**Figure S8** Principal component analysis (PCA) biplots for 106,427 variants from the one-million-variant clustering subset of the wild *Brassica oleracea* population. Variant annotation variables are shown as loading vectors. Panels **A – C** show variants coloured according to variant quality class based on the three-component Gaussian mixture model clustering. Green = high, purple = intermediate, orange = low.

**Table S4** Summary statistics for SNP quality classes based on GMM clustering in the wild *Brassica oleracea* population

| SNP Set | Variant Class | #SNPs | #True SNPs* | *Med(x)* | | | | | |
|---|---|---|---|---|---|---|---|---|---|
| | | | | QD | FS | SOR | MQ | MQRS | RPRS |
| Clustering Subset** (*k = 3*) | High | 226,925 | 395 | 13.4 | 2.4 | 0.7 | 60.0 | 0 | 0.1 |
| | Intermediate | 406,688 | 80 | 11.6 | 2.0 | 0.7 | 53.8 | -0.9 | 0.2 |
| | Low | 185,921 | 21 | 7.1 | 16.3 | 2.4 | 50.2 | -2.2 | 0.3 |
| 13k Truth Set | Pass | 13,310 | 13,310 | 15.6 | 2.0 | 0.7 | 60.0 | 0 | 0.2 |
| | Fail | 186 | 186 | 6.0 | 6.8 | 3.0 | 59.6 | 0 | 0.3 |

*k* number of clusters*; QD* QualByDepth*; FS* FisherStrand*; SOR* StrandOddsRatio*; MQ* MappingQuality*; MQRS* MappingQualityRankSum*; RPRS* ReadPosRankSum
*Variants overlapping with the 13k truth set
**Clustering Subset with SNPs containing only complete annotation profiles

#### Cohort-specific variant recalibration and tranche filtering

**Table S5** Summary statistics for cohort-specific SNP quality score recalibration across different truth sensitivity tranches in the wild *Brassica oleracea* population

| Truth Sensitivity Tranche | Known Sites | Novel Sites | Known Sites [Ti/Tv] | Novel Sites [Ti/Tv] | minVQSLod | Model | Accessible Truth Sites | Calls Truth Sites |
|---|---|---|---|---|---|---|---|---|
| 90.00 | 2,404 | 1,138,478 | 2.35 | 1.35 | 7.10 | SNP | 226,543 | 203,888 |
| 95.00 | 8,586 | 4,259,097 | 2.19 | 1.32 | 6.23 | SNP | 226,543 | 215,215 |
| 99.00 | 11,053 | 7,695,359 | 2.25 | 1.40 | 3.00 | SNP | 226,543 | 224,277 |
| 99.50 | 12,622 | 12,181,096 | 2.28 | 1.46 | 1.37 | SNP | 226,543 | 225,410 |
| 99.90 | 13,099 | 16,465,603 | 2.28 | 1.49 | -0.99 | SNP | 226,543 | 226,316 |
| 99.99 | 13,215 | 17,964,298 | 2.28 | 1.50 | -3.88 | SNP | 226,543 | 226,520 |
| 100.00 | 13,496 | 22,484,808 | 2.29 | 1.52 | -39855.10 | SNP | 226,543 | 226,543 |

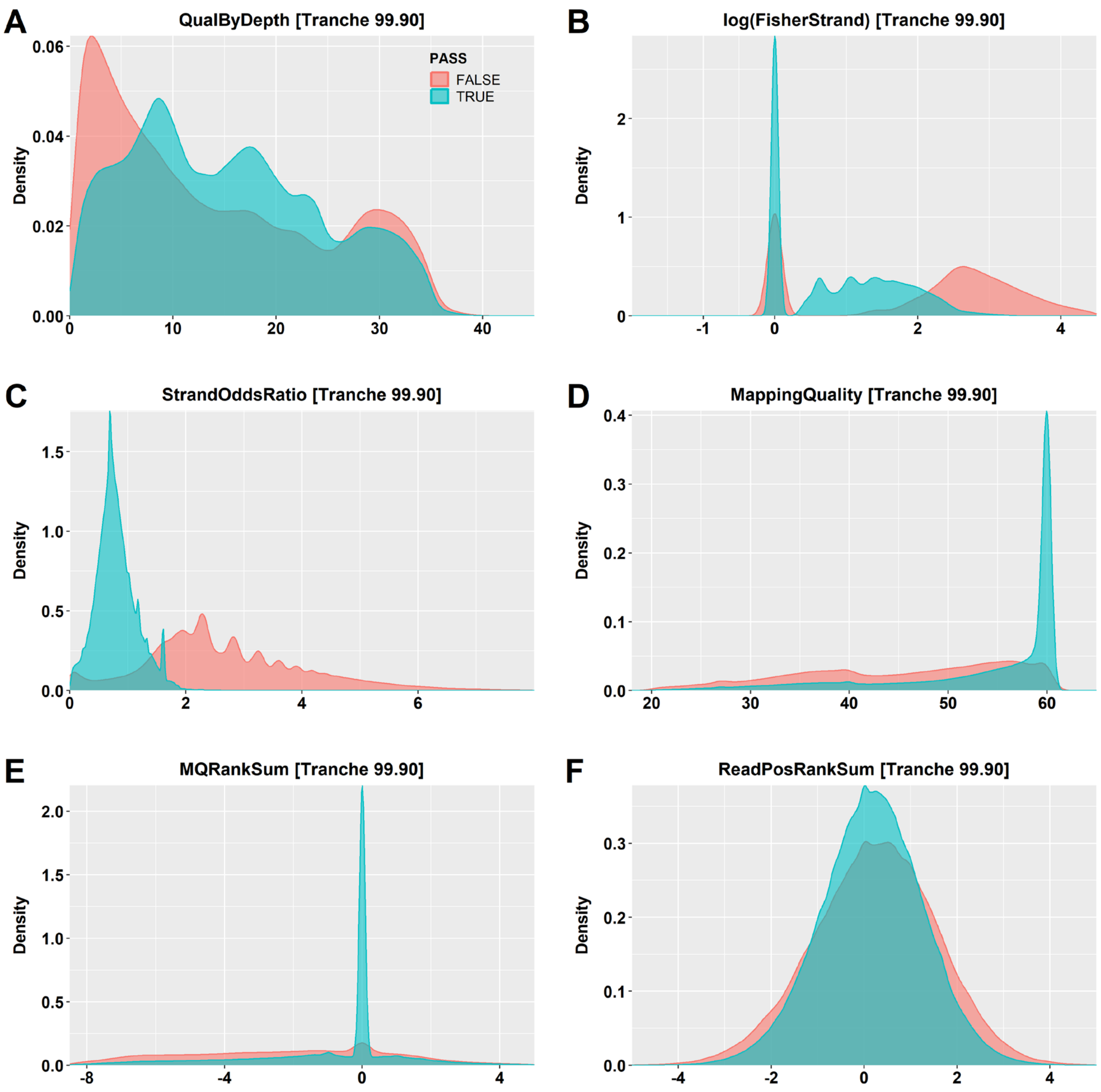


**Figure S8** Distribution of **A** QualByDepth, **B** log-transformed Fisher Strand, **C** Strand Odds Ratio, **D** Mapping Quality, **E** MQRankSum, and **F** ReadPosRankSum values for retained (blue) and filtered (red) SNPs following variant quality score recalibration (VQSR) and filtering at the 99.90% tranche in a one-million-variant subset from the wild *Brassica oleracea* population.
