## Supplementary Data 3 for "Unsupervised Clustering Enables Cohort-Specific Variant Recalibration in Crops"

### Unsupervised Clustering Enables Cohort-Specific Variant Recalibration in Crops

Thomas Bergmann[1,2], Tessa MacNish[1], Jacqueline Batley[2], and David Edwards[1]

[1]Centre for Applied Bioinformatics, University of Western Australia, 6009 Perth, Australia

[2]School of Biological Sciences, University of Western Australia, 6009 Perth, Australia

#### Variant Calling and Filtering in a *Triticum aestivum* Population

Scripts and resources:

https://github.com/tbergmann92/ Advanced-Crop-Variant-Discovery

#### Distribution of the SNP array-derived known variant sites in *Triticum aestivum*

**Table S1** Identification and distribution of 576,576 known SNP sites derived from the Wheat 660k SNP array across the *Triticum aestivum* 'AK58' reference genome

| Chromosome | Length [bp] | Known Sites | |
|---|---|---|---|
| | | SNP Count | bp between SNPs |
| 1A | 604,927,367 | 36,619 | 16,519 |
| 1B | 748,715,293 | 22,059 | 33,941 |
| 1D | 499,058,306 | 18,461 | 27,033 |
| 2A | 797,928,436 | 37,517 | 21,268 |
| 2B | 805,712,850 | 43,506 | 18,519 |
| 2D | 658,671,774 | 21,085 | 31,238 |
| 3A | 757,136,854 | 25,006 | 30,278 |
| 3B | 868,993,756 | 54,590 | 15,918 |
| 3D | 627,290,677 | 15,396 | 40,743 |
| 4A | 751,611,623 | 30,542 | 24,609 |
| 4B | 687,304,658 | 17,582 | 39,091 |
| 4D | 526,423,633 | 14,242 | 36,962 |
| 5A | 717,457,865 | 27,739 | 25,864 |
| 5B | 715,052,430 | 39,175 | 18,252 |
| 5D | 577,163,422 | 19,626 | 29,408 |
| 6A | 627,465,879 | 21,812 | 28,767 |
| 6B | 726,140,029 | 33,524 | 21,660 |
| 6D | 499,372,107 | 16,367 | 30,510 |
| 7A | 747,387,442 | 35,756 | 20,902 |
| 7B | 757,817,953 | 23,501 | 32,246 |
| 7D | 645,119,121 | 22,471 | 28,708 |
| | | **576,576** | **26,311** |

#### SNP array-derived variant sites improve base quality score recalibration in *T. aestivum*

1. Input data

- A recalibration file (csv) generated by *gatk AnalyzeCovariates*

2. Generation of bias plots before and after BQSR

Diagnostic plots for base quality scores, dinucleotide context, and machine cycle covariate bias are plotted with a custom R script:

- Plot_BQSR_Results_Wheat.r

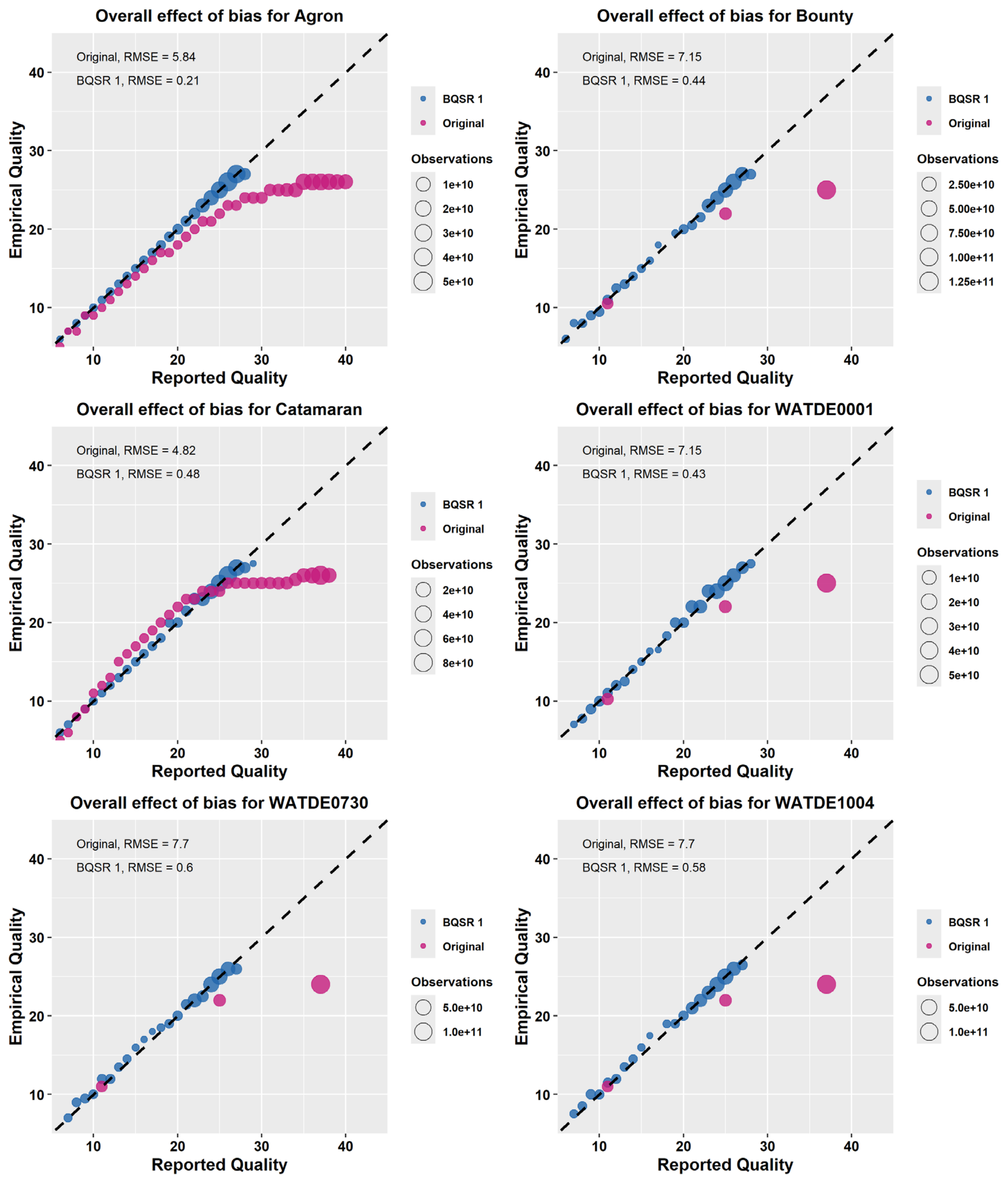


**Figure S1**.Base quality score recalibration (BQSR) with the 570k known sites in the *Triticum aestivum* 'AK58' reference genome for the base quality score bias across six samples from the wheat collection. The raw uncalibrated data contains un-binned and binned base quality scores: Q11, Q25, Q37. Pink = uncalibrated data; blue = recalibrated data.

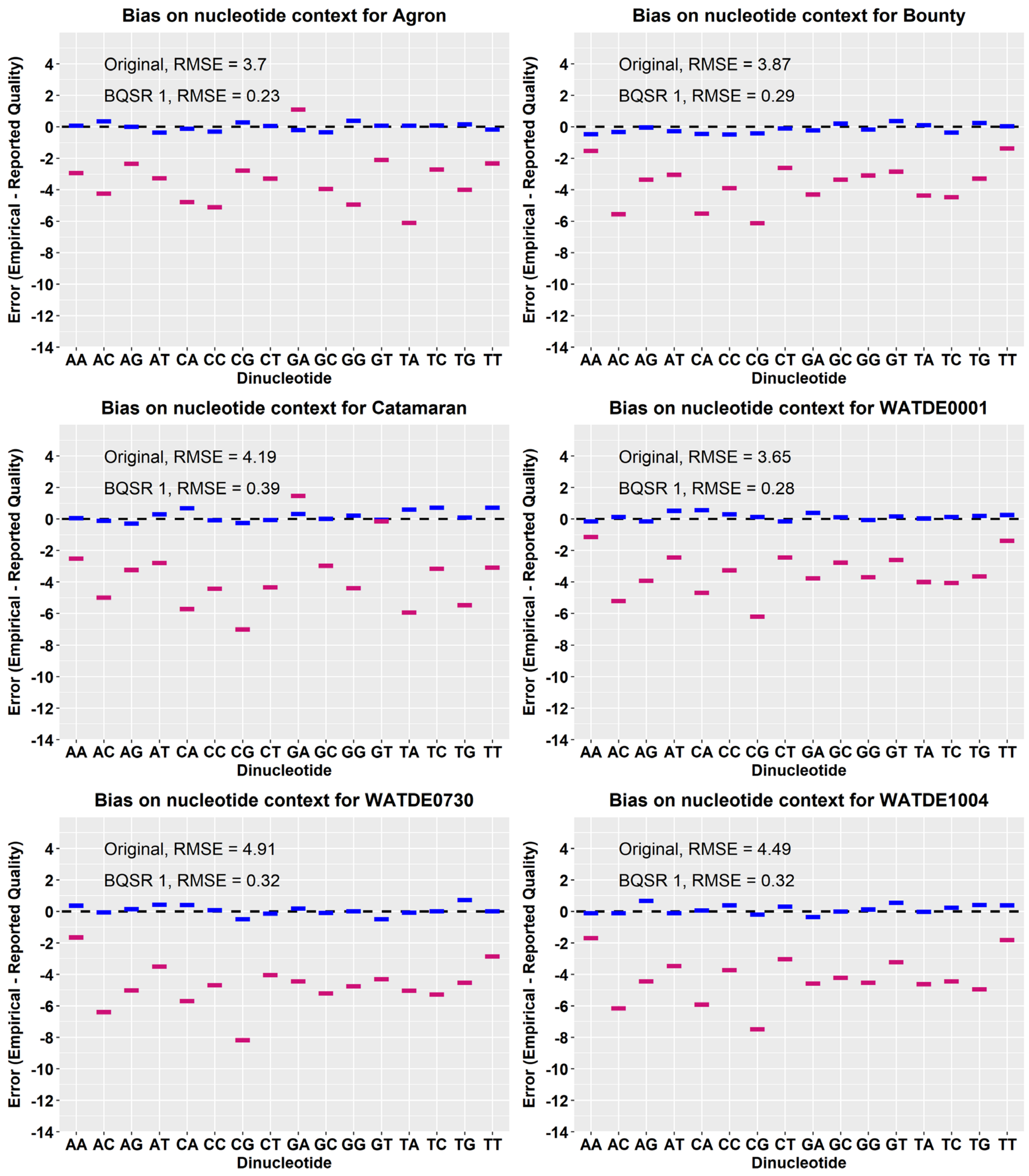


**Figure S2**.Base quality score recalibration (BQSR) with the 570k known sites in the *Triticum aestivum* ‘AK58’ reference genome for the dinucleotide context bias across six samples from the wheat collection. Pink = uncalibrated data; blue = recalibrated data

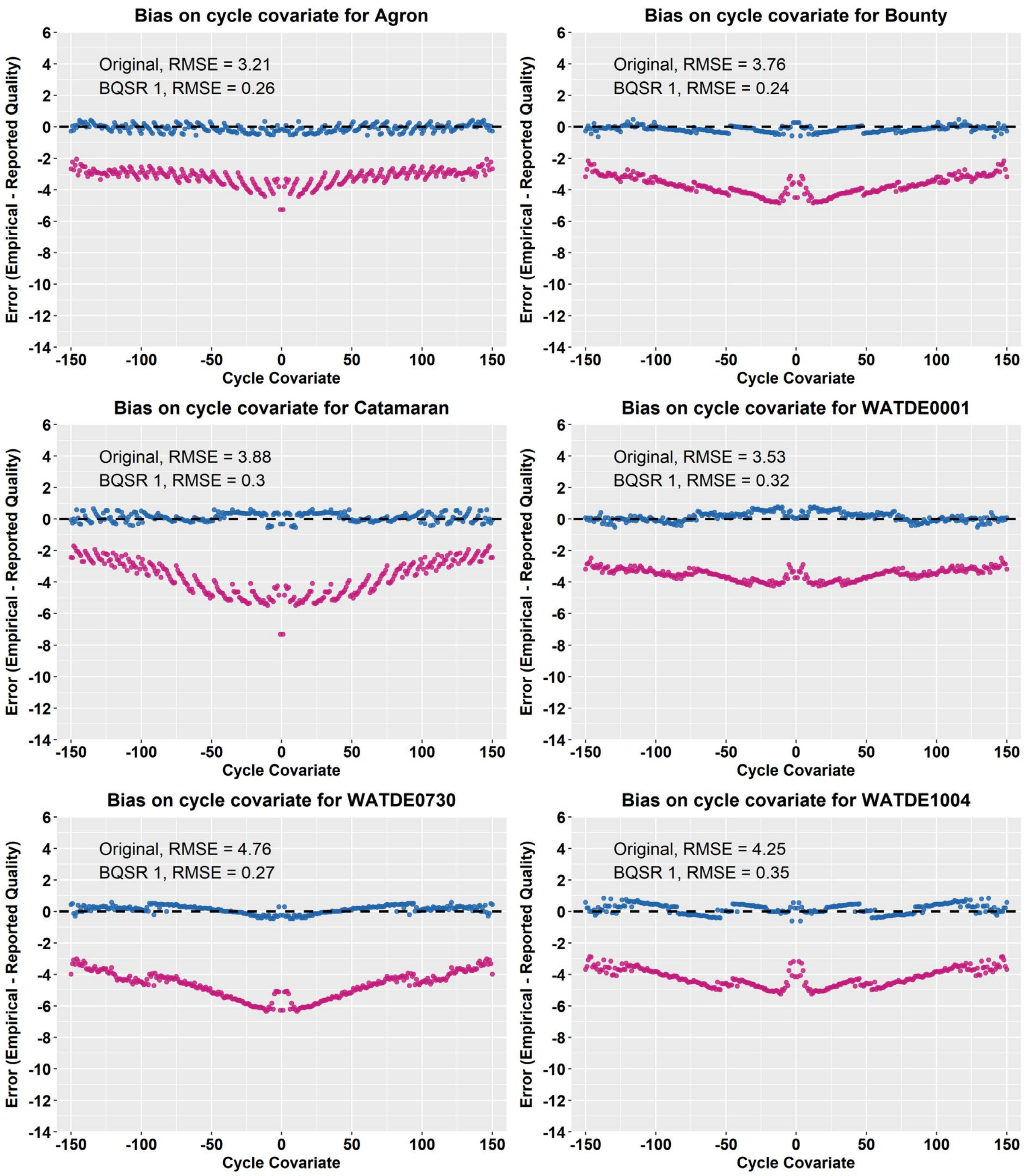


**Figure S3**.Base quality score recalibration (BQSR) with the 570k known sites in the *Triticum aestivum* ‘AK58’ reference genome for the machine cycle covariate bias across seven samples from the wheat collection. Pink = uncalibrated data; blue = recalibrated data.

#### Constructing a high-confidence truth set using SNP array-derived variant sites

After data pre-processing and the first round of raw variant calling (Table 2), matching variant sites between the unfiltered VCF and the 570k known sites are retrieved to create a cohort-specific truth set. This identifies 249,590 out of 71,561,408 SNP sites in the unfiltered VCF as true variant sites in the *Triticum aestivum* collection, hereafter referred to as 250k truth set.

**Table S2** Variant calling summary in the *Triticum aestivum* collection

| Feature | Raw VCF | 250k Truth Set |
|---|---|---|
| Total variants | 77,674,245 | 249,590 |
| SNP loci | 71,561,408 | 249,590 |
| INDEL loci | 5,992,962 | - |
| Ti/Tv-Ratio | 1.95 | 2.45 |

*Statistics were generated with gatk VariantEval*

> *Details on how to generate the truth set are found in Supplementary Data 1 in the corresponding section.*

#### Preparing a three-million-variant clustering subset for annotation analysis

To account for the large genome size of wheat, the AK58 genome was divided into 5 kb intervals, and a single variant site was randomly sampled from each interval. The code to generate this subset is provided below.

1. Input data

- The tab-delimited annotation table of the unfiltered VCF: *$RAW_ANNOTATION_TSV*

2. Creating the clustering subset:

**###**

**# Select a single SNP from each 5kb interval:**

```
awk '
BEGIN{
   FS=OFS="\t"
   srand()
}
$3=="SNP" {
   bin=$1":"int(($2-1)/5000)

   if (!(bin in n)) {
      n[bin]=1
      pick[bin]=$0
   } else {
      n[bin]++
      if (rand() < 1/n[bin])
         pick[bin]=$0
   }
}
END{
   for(i in pick)
      print pick[i]
}' $RAW_ANNOTATION_TSV | sort -k1,1 -k2,2n > $SNP_SUBSET_ANNOTATIONS_TSV
```

**###**

**Table S3** Summary of SNP distribution in the three-million-variant clustering subset of the *Triticum aestivum* collection

| Chromosome | Length [bp] | Known Sites | |
|---|---|---|---|
| | | SNP Count | bp between SNPs |
| 1A | 604,927,367 | 117,173 | 5,163 |
| 1B | 748,715,293 | 99,556 | 7,521 |
| 1D | 499,058,306 | 95,752 | 5,212 |
| 2A | 797,928,436 | 153,638 | 5,194 |
| 2B | 805,712,850 | 155,584 | 5,179 |
| 2D | 658,671,774 | 126,577 | 5,204 |
| 3A | 757,136,854 | 144,612 | 5,236 |
| 3B | 868,993,756 | 167,877 | 5,176 |
| 3D | 627,290,677 | 119,817 | 5,235 |
| 4A | 751,611,623 | 144,766 | 5,192 |
| 4B | 687,304,658 | 132,805 | 5,175 |
| 4D | 526,423,633 | 101,078 | 5,208 |
| 5A | 717,457,865 | 138,508 | 5,180 |
| 5B | 715,052,430 | 138,105 | 5,178 |
| 5D | 577,163,422 | 109,840 | 5,255 |
| 6A | 627,465,879 | 121,454 | 5,166 |
| 6B | 726,140,029 | 140,691 | 5,161 |
| 6D | 499,372,107 | 95,253 | 5,243 |
| 7A | 747,387,442 | 144,382 | 5,176 |
| 7B | 757,817,953 | 147,027 | 5,154 |
| 7D | 645,119,121 | 123,476 | 5,225 |
| | | **2,717,971** | **5,288** |

#### Comparing variant annotation profiles between call sets

*~.Details on how to analyse the variant context annotations are found in Supplementary Data 1 in the corresponding section.*

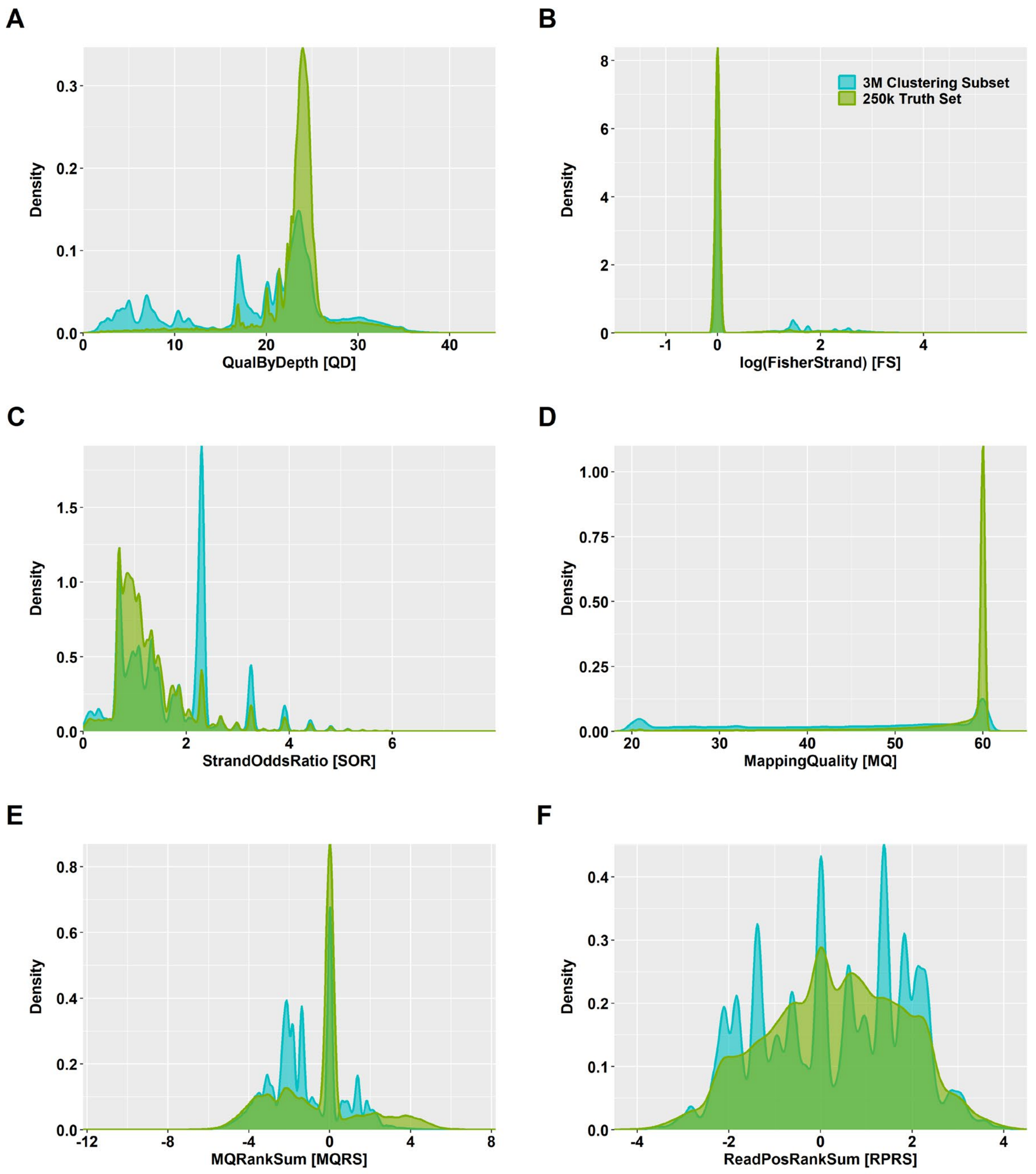


**Figure S4** Density plots for **A** QualByDepth (QD), **B** Fisher Strand (FS), **C** Strand Odds Ratio (SOR), **D** Mapping Quality (MQ), **E** MQRankSum (MQRS), and **F** ReadPosRankSum (RPRS) in the three-million-variant clustering subset (blue) and the 250k truth set (green) in the *Triticum aestivum* population.

#### Characterization of the multi-dimensional annotation space for clustering

*~. Details on how to analyse the multi-dimensional annotation space are found in Supplementary Data 1 in the corresponding section.*

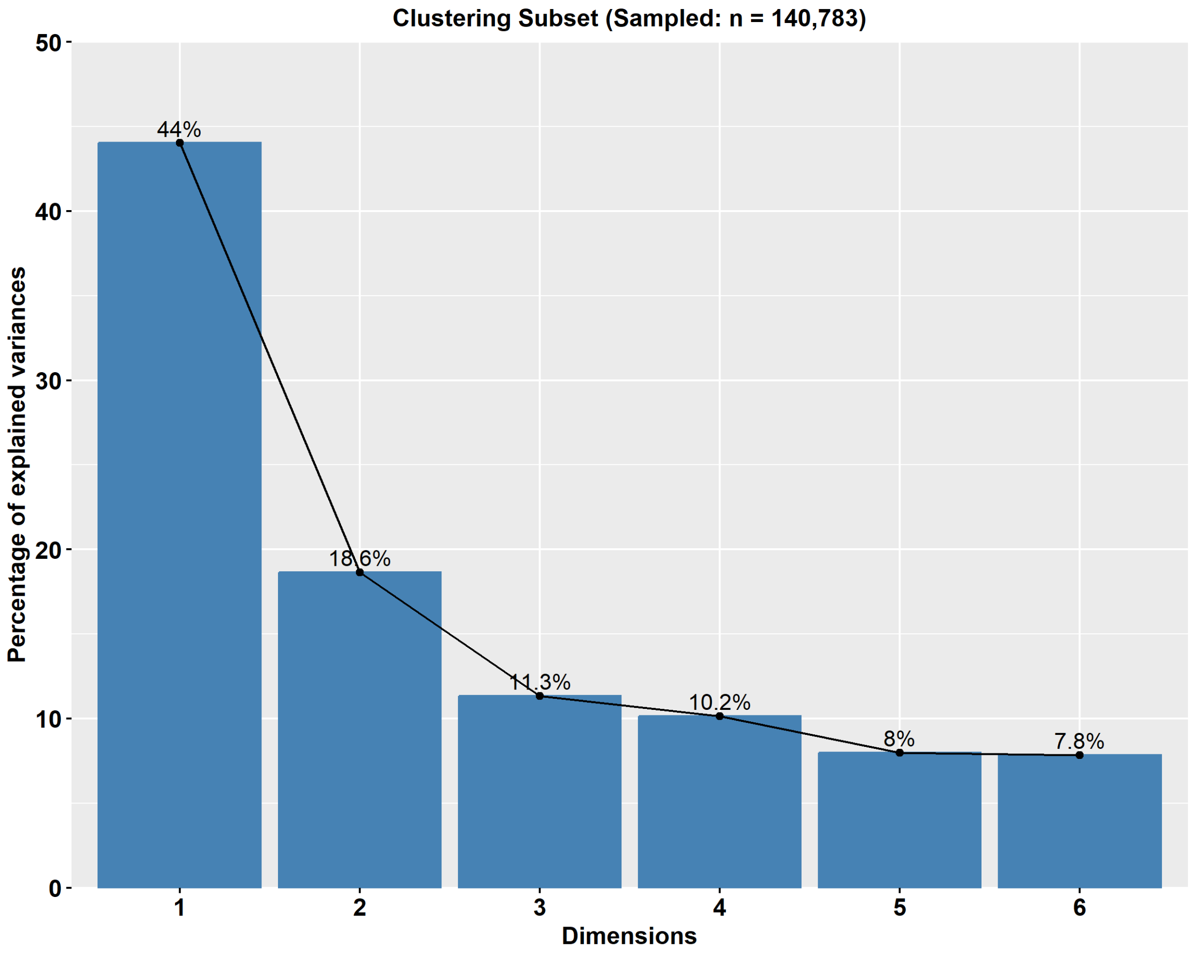


**Figure S5** Scree plots of principal components for six variant context annotations in the three-million-variant clustering subset from the *Triticum aestivum* population. The principal component analysis was performed with complete observations for QualByDepth, Fisher Strand, Strand Odds Ratio, Mapping Quality, MQRankSum, and ReadPosRankSum.

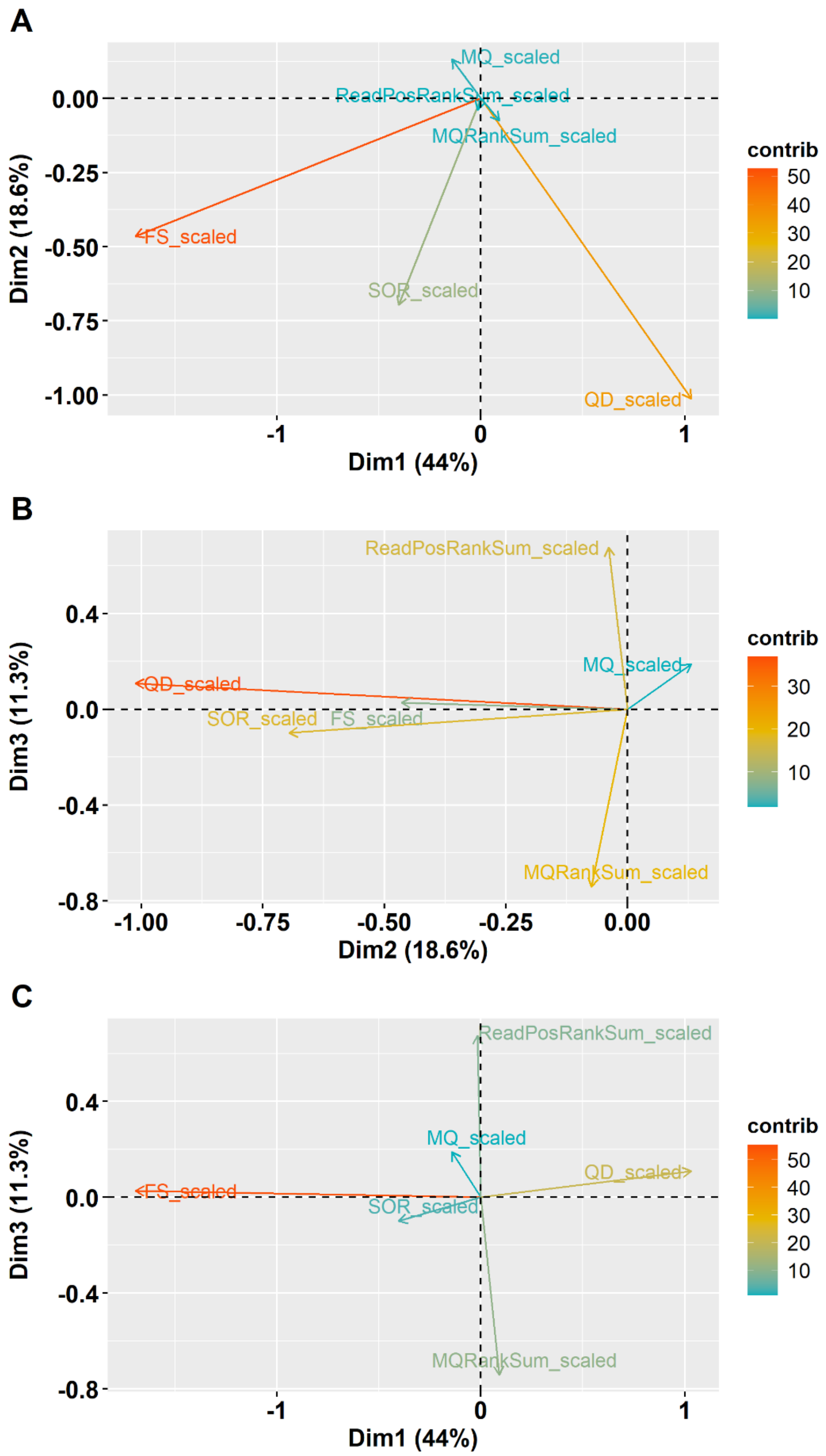


**Figure S6**.Loading plots of the principal components (PCs) based on ~150k SNPs from the three-million-variant clustering subset of the *Triticum aestivum* population. **A – C** Loading plots for PCs 1 – 3 with six variant context annotations. *QD* QualByDepth *FS* FisherStrand *SOR* StrandOddsRatio *MQ* MappingQuality

#### Unsupervised GMM clustering identifies high-quality variant classes

*> Details on how to build the model are found in Supplementary Data 1 in the corresponding section.*

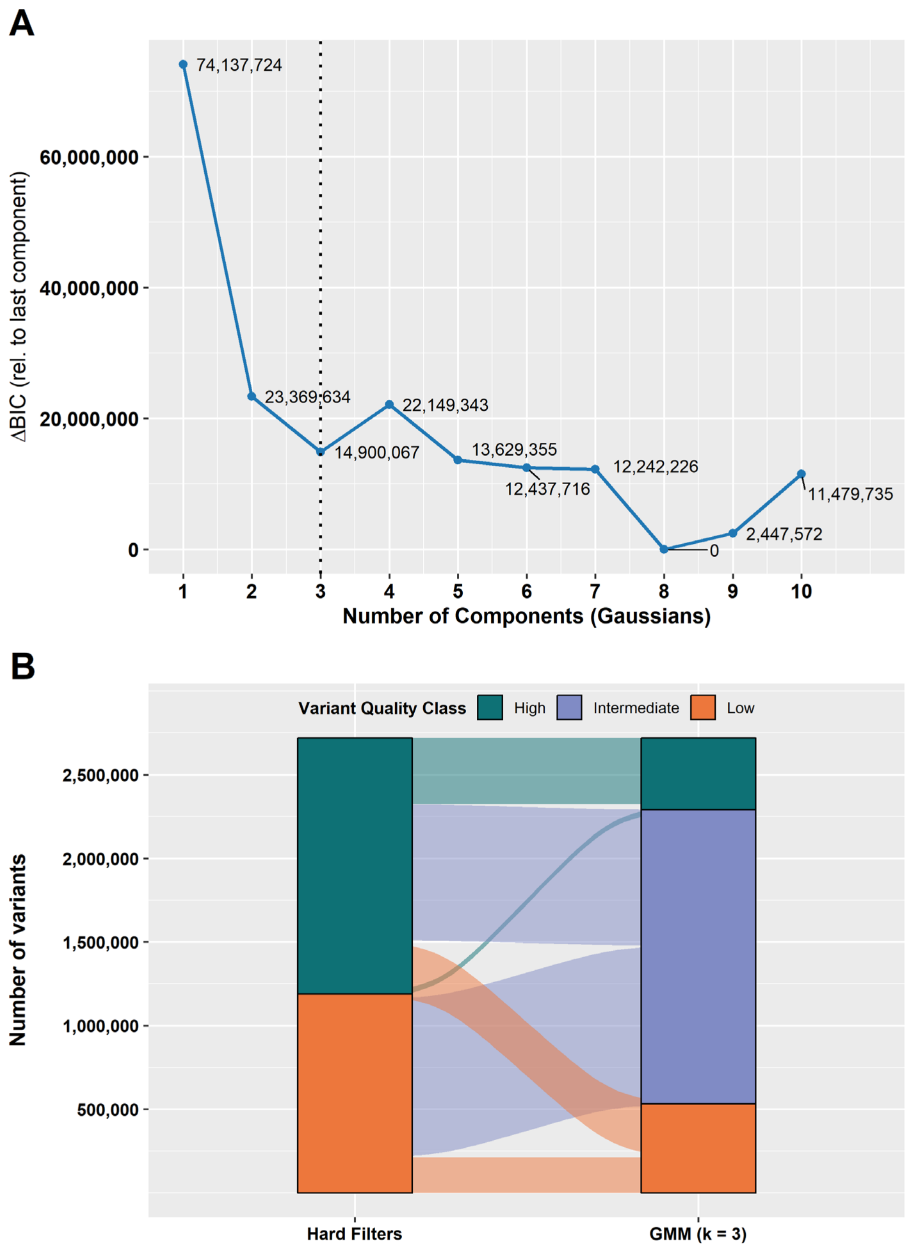


**Figure S7** Model selection and variant quality class stratification using Gaussian mixture model (GMM)-based clustering. **A** Change in Bayesian Information Criterion (ΔBIC) relative to the largest model ($k$ = 10) across increasing numbers of Gaussian components ($k$). Lower values indicate better model support. The dashed vertical line indicates best-model-fit. **B** Alluvial plot showing the flow of variant classification between hard filtering and GMM-based clustering ($k$ = 3). Each stratum represents the classification at a given stage, and its vertical width corresponds to the number of variants. Alluvia are coloured according to group assignments from the GMM with $k$ = 3.

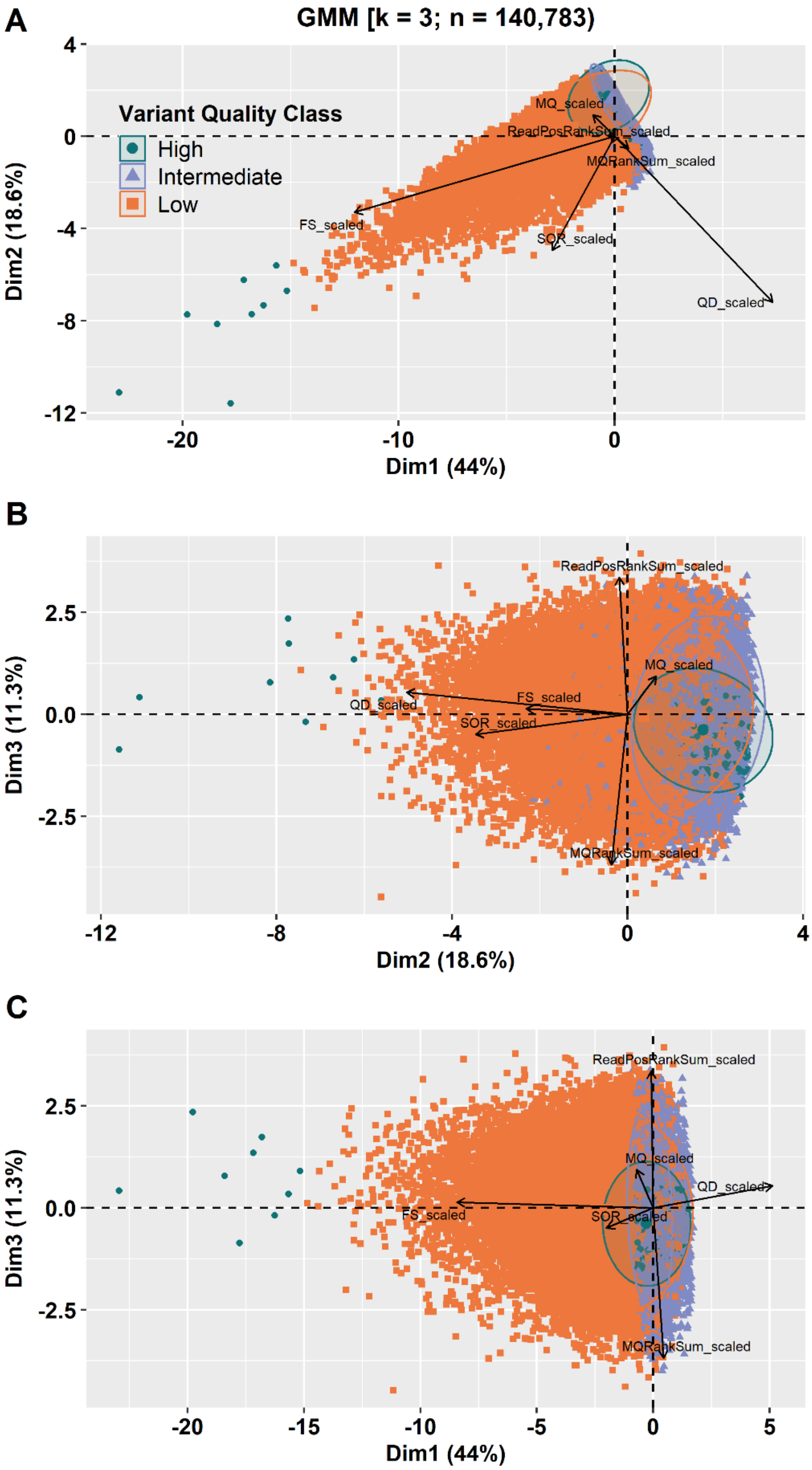


**Figure S8** Principal component analysis (PCA) biplots for 140,783 variants from the three-million-variant clustering subset of the *Triticum aestivum* population. Variant annotation variables are shown as loading vectors. Panels **A – C** show variants coloured according to variant quality class based on the three-component Gaussian mixture model clustering. Green = high, purple = intermediate, orange = low.

**Table S4** Summary statistics for SNP quality classes based on GMM clustering in the *Triticum aestivum* population

| SNP Set | Variant Class | #SNPs | #True SNPs* | *Med(x)* | | | | | |
|---|---|---|---|---|---|---|---|---|---|
| | | | | QD | FS | SOR | MQ | MQRS | RPRS |
| Clustering Subset** (*k* = *3*) | High | 427,767 | 4,456 | 23.1 | 0 | 1.3 | 60 | 0 | 0.1 |
| | Intermediate | 1,759,024 | 2,904 | 22.9 | 0 | 1.3 | 40 | -1.4 | 0.5 |
| | Low | 531,176 | 401 | 7.4 | 4.8 | 2.2 | 49 | -1.4 | 0.5 |
| 250k Truth Set | Pass | 222,379 | 222,379 | 23.9 | 0 | 1.1 | 60 | 0 | 0.3 |
| | Fail | 27,211 | 27,211 | 22.9 | 0 | 1.9 | 36 | -1.5 | 0.3 |

*k* number of clusters; *QD* QualByDepth; *FS* FisherStrand; *SOR* StrandOddsRatio; *MQ* MappingQuality; *MQRS* MappingQualityRankSum; *RPRS* ReadPosRankSum
*Variants overlapping with the 250k truth set
**Clustering Subset with SNPs containing only complete annotation profiles for QD, FS, SOR, and MQ

#### Cohort-specific variant recalibration and tranche filtering

**Table S5** Summary statistics for cohort-specific SNP quality score recalibration across different truth sensitivity tranches in the *Triticum aestivum* population

| Truth Sensitivity Tranche | Known Sites | Novel Sites | Known Sites [Ti/Tv] | Novel Sites [Ti/Tv] | minVQSLod | Model | Accessible Truth Sites | Calls Truth Sites |
|---|---|---|---|---|---|---|---|---|
| 90.00 | 94,718 | 7,456,632 | 2.50 | 2.27 | 4.06 | SNP | 427,696 | 384,926 |
| 95.00 | 131,784 | 14,519,821 | 2.50 | 2.09 | 3.41 | SNP | 427,696 | 406,311 |
| 99.00 | 194,150 | 37,027,606 | 2.48 | 2.08 | 1.89 | SNP | 427,696 | 423,419 |
| 99.50 | 207,152 | 43,730,236 | 2.48 | 2.10 | 1.28 | SNP | 427,696 | 425,557 |
| 99.90 | 219,055 | 50,970,949 | 2.48 | 2.11 | 0.19 | SNP | 427,696 | 427,268 |
| 99.99 | 225,502 | 56,044,975 | 2.48 | 2.12 | -1.15 | SNP | 427,696 | 427,653 |
| 100.00 | 249,173 | 71,312,235 | 2.45 | 1.95 | -10067511.26 | SNP | 427,696 | 427,696 |

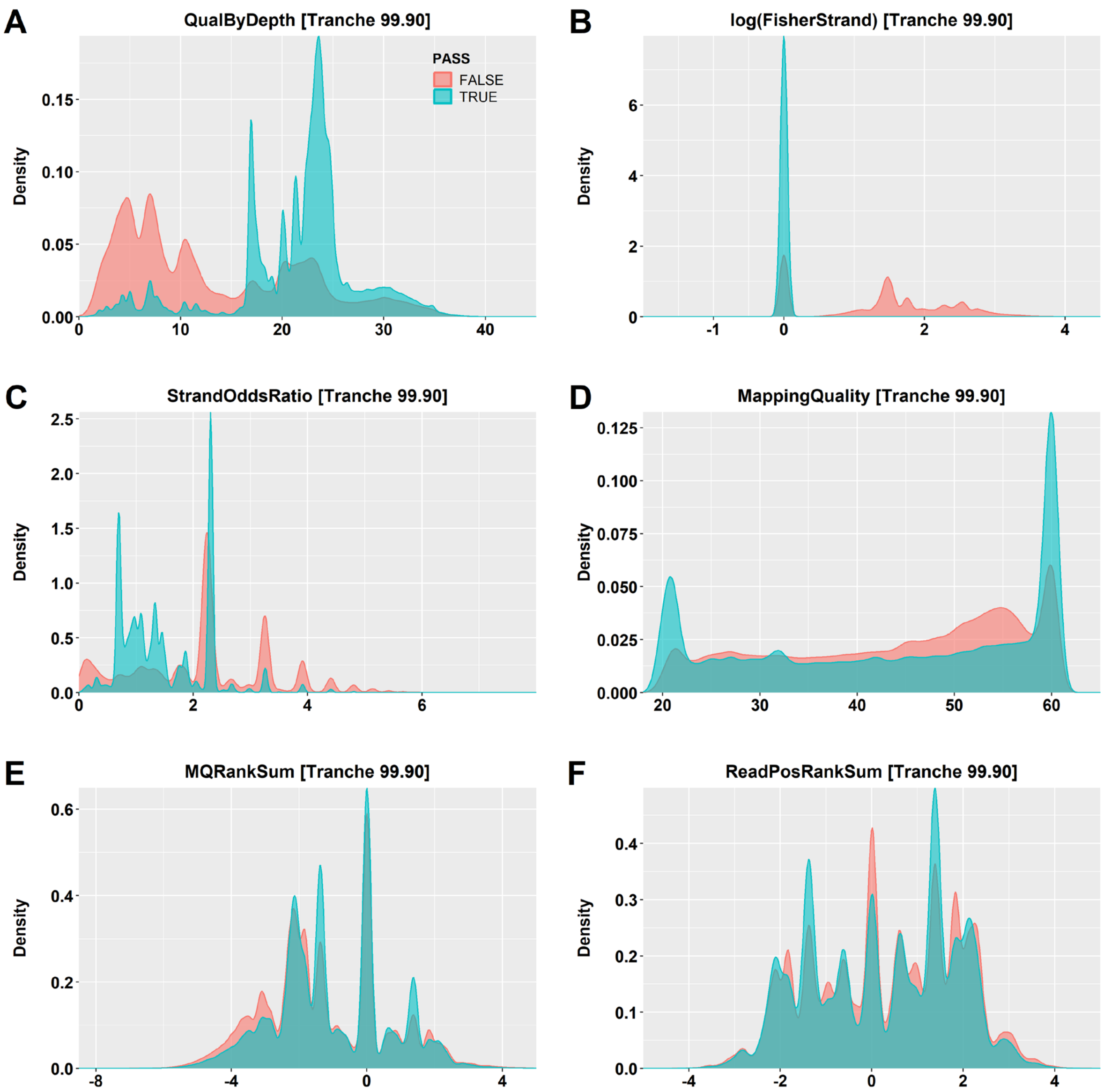


**Figure S9** Distribution of **A** QualByDepth, **B** log-transformed Fisher Strand, **C** Strand Odds Ratio, **D** Mapping Quality, **E** MQRankSum, and **F** ReadPosRankSum values for retained (blue) and filtered (red) SNPs following variant quality score recalibration (VQSR) and filtering at the 99.90% tranche in a three-million-variant subset from the *Triticum aestivum* population.
